# The Genome of the Charophyte Alga *Penium margaritaceum* Bears Footprints of the Evolutionary Origins of Land Plants

**DOI:** 10.1101/835561

**Authors:** Chen Jiao, Iben Sørensen, Xuepeng Sun, Honghe Sun, Hila Behar, Saleh Alseekh, Glenn Philippe, Kattia Palacio Lopez, Li Sun, Reagan Reed, Susan Jeon, Reiko Kiyonami, Sheng Zhang, Alisdair R. Fernie, Harry Brumer, David S. Domozych, Zhangjun Fei, Jocelyn K. C. Rose

**Author notes:** These authors contributed equally to this work (Chen Jiao, Iben Sørensen, Xuepeng Sun). Correspondence should be addressed to Jocelyn K. C. Rose, Zhangjun Fei and David S. Domozych.

## Abstract

The colonization of land by plants was a pivotal event in the history of the biosphere, and yet the underlying evolutionary features and innovations of the first land plant ancestors are not well understood. Here we present the genome sequence of the unicellular alga *Penium margaritaceum*, a member of the Zygnematophyceae, the sister lineage to land plants. The *P. margaritaceum* genome has a high proportion of repeat sequences, which are associated with massive segmental gene duplications, likely facilitating neofunctionalization. Compared with earlier diverging plant lineages, *P. margaritaceum* has uniquely expanded repertoires of gene families, signaling networks and adaptive responses, supporting its phylogenetic placement and highlighting the evolutionary trajectory towards terrestrialization. These encompass a broad range of physiological processes and cellular structures, such as large families of extracellular polymer biosynthetic and modifying enzymes involved in cell wall assembly and remodeling. Transcriptome profiling of cells exposed to conditions that are common in terrestrial habitats, namely high light and desiccation, further elucidated key adaptations to the semi-aquatic ecosystems that are home to the Zygnematophyceae. Such habitats, in which a simpler body plan would be advantageous, likely provided the evolutionary crucible in which selective pressures shaped the transition to land. Earlier diverging charophyte lineages that are characterized by more complex land plant-like anatomies have either remained exclusively aquatic, or developed alternative life styles that allow periods of desiccation.

## INTRODUCTION

One of the most momentous evolutionary events in the history of life on Earth is thought to have occurred approximately 500 million years ago (Mya), when a single lineage of freshwater algae developed the capacity to colonize land (Delwiche and Timme, 2011). These pioneering oxygenating auxotrophs had a profound effect on the atmosphere and geochemical composition of the soil (Rensing, 2018; Delwiche and Cooper, 2015) and paved the way for an explosion of land plant diversification, and the evolution of other branches of terrestrial life. A key question in understanding the origins of life on land is the nature of the adaptive traits that enabled this remarkable transition.

Green plants (Viridiplantae) are comprised of the chlorophyte algae and the monophyletic group Streptophyta, comprising land plants (embryophytes) and the charophyte algae (Fig. 1A). Two groups of charophyte lineages have been defined: the earlier diverging Mesostigmatophyceae together with Chlorokybophyceae and Klebsormidophyceae; and the later diverging Charophyceae, Coleochaetophyceae and Zygnematophyceae. There is now considerable molecular evidence that the Zygnematophyceae are the closest relatives to land plants (Delwiche and Cooper, 2015; Wodniok et al., 2011; Leebens-Mack et al., 2019). This may be considered somewhat paradoxical, in that members of the Zygnematophyceae exhibit a notably simpler body plan (e.g. unicells, filaments) than taxa of the Charaophyceae or Coleochaetophyceae (e.g. complex branched filamentous aggregates, pseudoparenchymatous forms) and also undergo sexual reproduction via conjugation rather than oogamy. However, it has been postulated that these less complex characteristics of the Zygnematophyceae represent a manifestation of a reduction trend in their evolutionary history (Delwiche and Timme, 2011). The smaller and simpler growth habit, with an increased capacity to tolerate water stress, may have been advantageous to life in shallow, ephemeral wetlands, i.e., habitats that may have been common during the time of algal emergence onto land.

**Figure 1.**
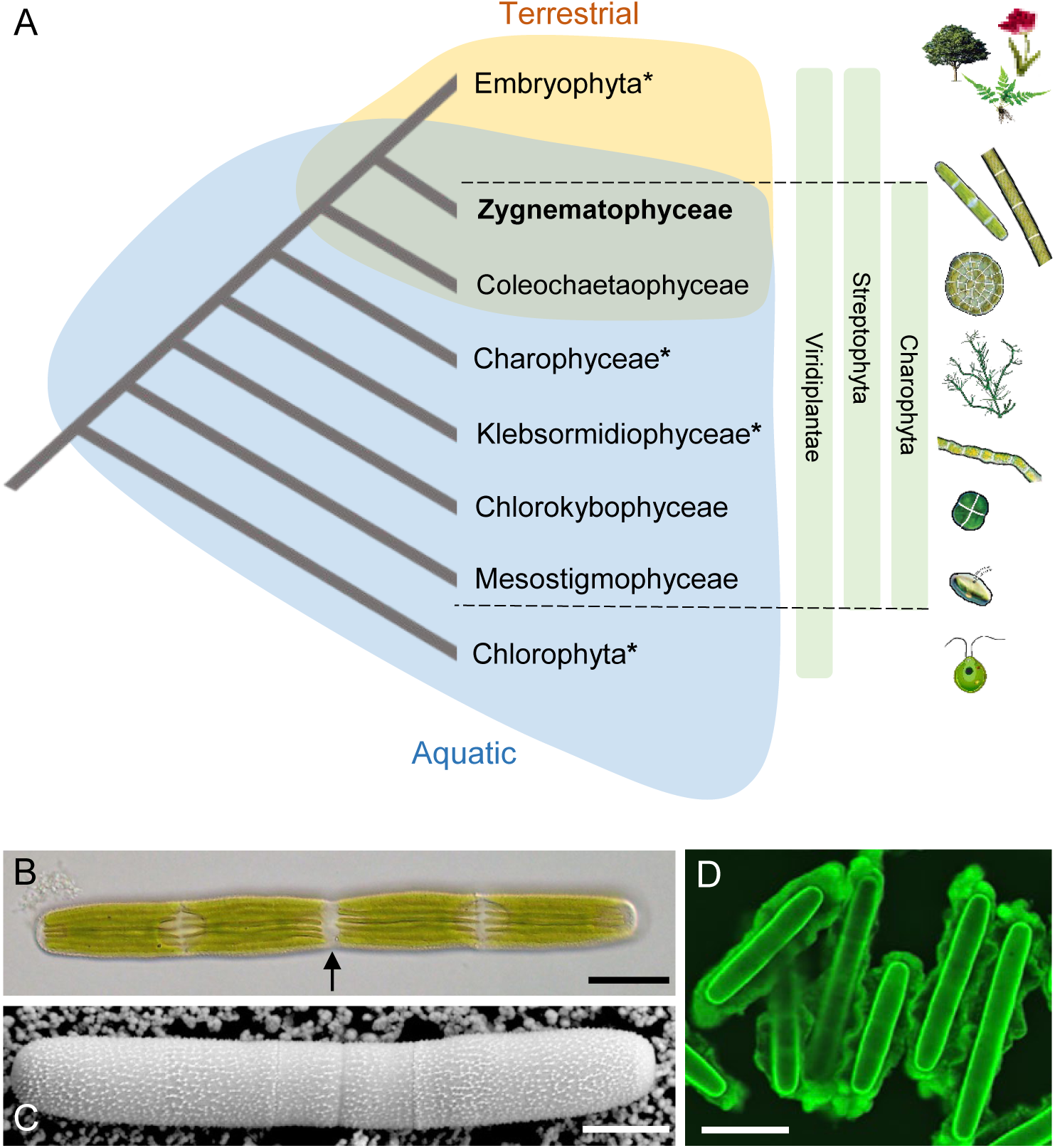
Phylogeny of green plants and morphology of *Penium margaritaceum*. (A) Current positioning of charophytes in plant phylogeny, highlighting variation in typical body plans (e.g. unicellular, filamentous and complex branched and multicellular) and associated terrestrial or aquatic habitats. Lineages for which there is a representative genome sequence are shown with an asterisk. (B) Cylindrical *P. margaritaceum* cell consisting of two semi-cells, each with one or two chloroplasts. The cell center (isthmus; arrow) contains the nucleus and is the major site of cell expansion. (C) Scanning electron micrograph image of a *P. margaritaceum* cell, highlighting the complex lattice of cell wall pectin polysaccharides on the cell surface. (D). Confocal scanning laser microscopy image of *P. margaritaceum* cells labeled with an antibody to the mucilage that encases them. Scale bars: B, 15 µm; C, 12 µm; D, 15 µm.

Extant charophytes have a remarkable range of body plans, even though fossil evidence suggests that they represent only a small proportion of the diversity that previously existed (Feist et al., 2005). This morphological diversity and its underlying developmental machinery provide a rich set of opportunities to elucidate adaptations that may have been critical for the invasion of land. Insights into the evolutionary origins and adaptive traits of taxa have been gleaned through analysis of the only two available genome sequences in the six Charophyte orders, Klebsormidophyceae (*Klebsormidium nitens;* Hori et al., 2014) and Charophyceae (*Chara braunii*; Nishiyama et al., 2018). In the earlier diverging *K. nitens*, the ability to synthesize certain phytohormones and associated signaling intermediates, along with a mechanism to cope with high light, represent physiological adaptations that were likely key for terrestrialization. The genome sequence of the later diverging *C. braunii*, revealed further innovations, including a reactive oxygen species response (ROS) network, the production of stress and storage proteins, components of canonical phytohormone biosynthetic and response pathways and elaboration of transcription factors for oogamous gamete production. However, a critical gap in the genomic information required to elucidate the evolutionary arc from aquatic to terrestrial plant life has been a well-defined genome representing the Zygnematophyceae, the sister group to embryophytes.

Here, we present the genome and transcriptomes of the unicellular desmid *Penium margaritaceum,* an archetype of the Zygnematophyceae with the simplest of body plans (Fig. 1B). *P. margaritaceum* has an architecturally complex land plant-like cell wall (Domozych et al., 2007; 2014; Sørensen et al., 2011) and secretes mucilaginous polysaccharides (Fig. 1C, D), which may be associated with its adaptation to living in transient wetlands that experience frequent drying. In this study, we sought to identify suites of genes that facilitate adaptation to an ephemeral semi-terrestrial life style, and to determine whether the relatively simple morphology of *P. margaritaceum* is associated with genome features, such as reductionism compared with other charophyte lineages that have more complex multicellular body plans. We further investigated the responses of *P. margaritaceum* to a range of abiotic stresses, in order to elucidate the adaptations that enable tolerance of the environmental challenges imposed by terrestrial habitats.

## RESULTS

### Genome and gene set of *P. margaritaceum*

We generated a total of 954 Gb Illumina paired-end and 433 Gb mate-pair sequences (Supplementary Table 1), representing 201× and 92× coverage, respectively, of the haploid genome of *P. margaritaceum* with an estimated size of 4.7 Gb (Supplementary Fig. 1A). Assembly of these sequences resulted in 332,786 scaffolds, with a cumulative size of 3.661 Gb and an N50 of 116.1 kb. The nuclear assembly captured most of the k-mers in the Illumina reads and low frequency k-mers representing sequencing errors were absent (Supplementary Fig. 1B). In addition, the mapping rates of genomic and RNA-Seq reads against the nuclear assembly were 97.5% and 96.8%, respectively (Supplementary Table 2). The single nucleotide polymorphism (SNP) frequency distribution on the 100 longest scaffolds was consistent with a haploid genome (Supplementary Fig. 1C). The mitochondrial and chloroplast genomes were also fully assembled, and comprised 95,332 and 145,411 nucleotides, respectively (Supplementary Fig. 2).

The assembly contains a large proportion (80.6%) of repeat sequences (Supplementary Table 3), particularly long terminal repeat (LTR) retrotransposons and simple repeats (Fig. 2A). Unlike land plants and *C*. *braunii*, in which *gypsy* is the predominant LTR family, the *P. margaritaceum* genome has a large proportion of *copia* retrotransposons, which are rare in other green algae and absent from *C*. *braunii* (Nishiyama et al. 2018). An estimation of divergence time indicated that the *copia* expansion in the *P. margaritaceum* genome was relatively recent, around 2.1 Mya (Fig. 2B). Retrotransposons carrying tyrosine recombinases, such as the DIRS and Ngaro families, which are found in some chlorophytes and both *C*. *braunii* and *K. nitens* genomes, are not present in *P. margaritaceum* and land plants (Fig. 2A).

**Figure 2.**
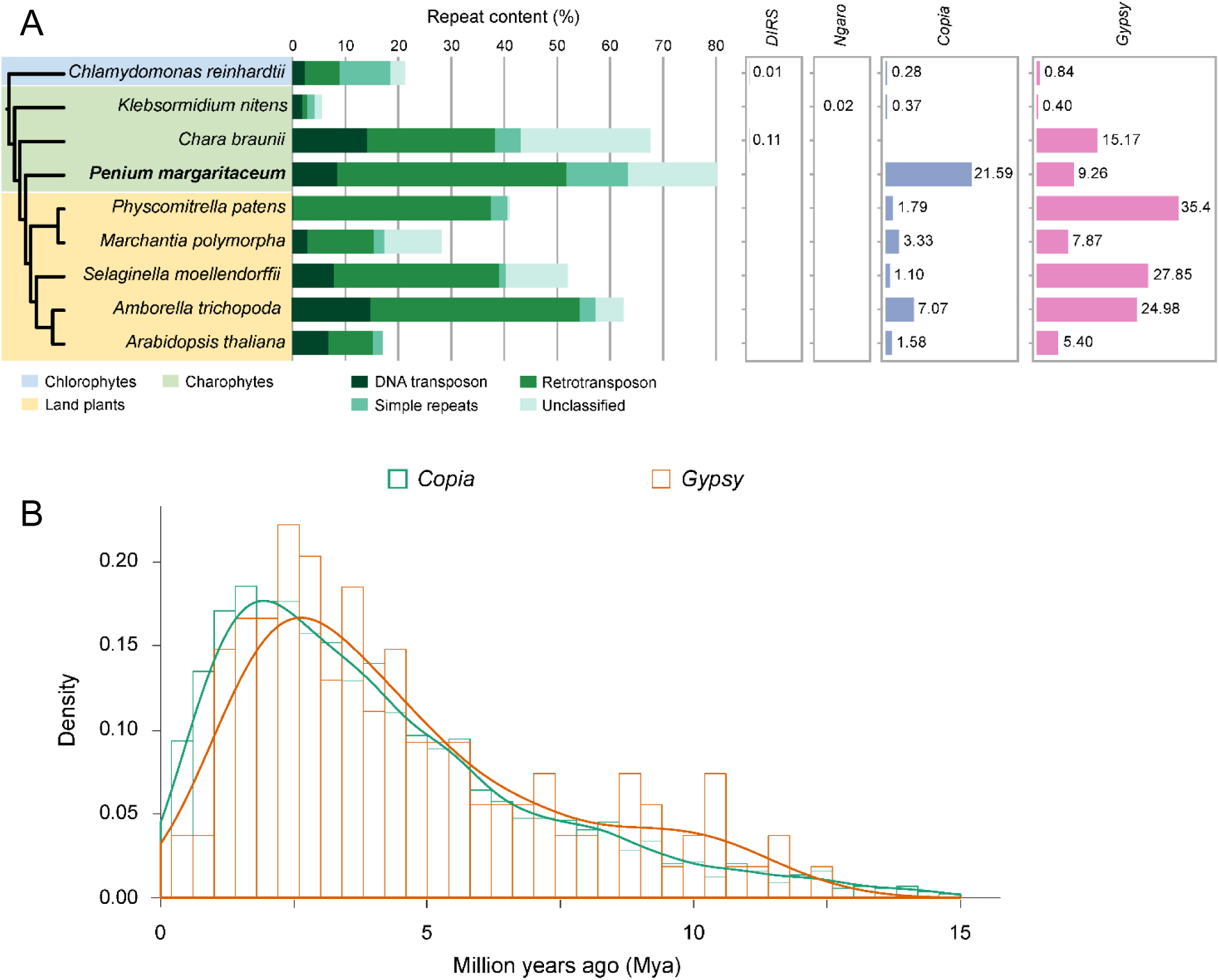
Repeat sequences in genomes of *P. margaritaceum* and selected green plant species. (A) Contents of different repeats in different green plant species. The schematic tree on the left shows evolutionary relationships. Numbers on the right panel correspond to *DIRS*, *Ngaro*, *copia*, and *gypsy* percentages. (B) Estimated insertion time of full-length *copia* and *gypsy* LTRs in the *P. margaritaceum* genome (band width: 0.4).

We predicted 52,333 high-confidence protein-coding genes in the *P. margaritaceum* genome, of which 99.3% were supported either by Illumina RNA-Seq data, or by homologs in the NCBI non-redundant protein database. Assessment of gene space completeness using BUSCO (Simão et al., 2015) indicated a low rate of missing genes (8.25%), but the fragmented gene rate was relatively high (21.45%). To inform the annotation, we performed transcriptome sequencing with PacBio Iso-Seq technology, which generated 52,134 full-length transcripts consisting of 73,813 isoforms. These PacBio transcripts, together with 145,267 representative transcripts assembled from Illumina RNA-Seq data, were used to build a master gene set by integrating with the high-confidence gene models. The final master gene set consisted of 53,262 genes (47,863 from the high-confidence gene models, 2,391 from the PacBio transcripts and 3,008 from the Illumina transcript data). The missing and fragmented gene rates of the master gene set were 2.31% and 13.20%, respectively, and the complete BUSCO rate was 84.49% (Supplementary Table 4).

### Genome and gene family evolution

Orthologous genes among *P. margaritaceum* and 13 representative species spanning the green plant lineage were identified. Phylogenetic analysis of low-copy orthologous groups confirmed the close relationship between *P. margaritaceum* and land plants, and indicated that *P. margaritaceum* diverged from the common ancestor of land plants around 663-552 Mya (Fig. 3A). This Proterozoic separation substantially predates a proposed crown origin of embryophytes 492-461 Mya in the Phanerozoic (middle Cambrian-early Ordovician), but is consistent with a recent estimate (Morris et al., 2108). The *P. margaritaceum* genome has not undergone any whole genome duplication (WGD) events (Supplementary Fig. 3), unlike the multiple rounds that have occurred in land plants (Van de Peer et al., 2017). However, substantial segmental gene duplications were found in the *P. margaritaceum* genome, which is consistent with the high TE abundance, given that massive segmental gene duplications are often found in organisms with a high TE content (Panchy et al., 2016).

**Figure 3.**
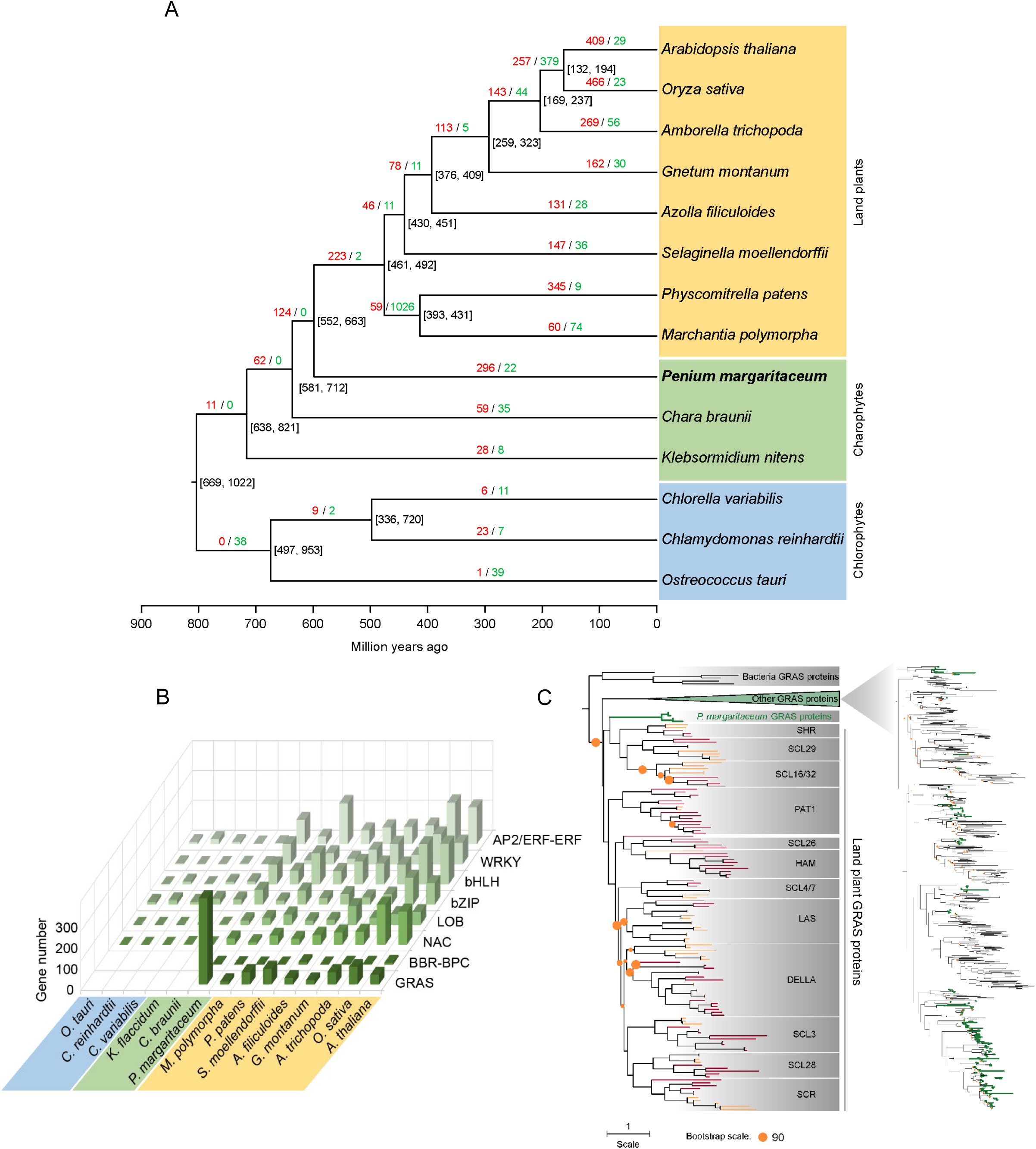
Gene family evolution in green plant species. (A) Divergence and gene family evolution of selected species. Numbers in bracket indicate the estimated age in million years. Numbers on branches represent significantly (P < 0.05) expanded (red) or contracted (green) gene families in that branch compared to its last ancestor. (B) Transcription factor families expanded compared to earlier diverging lineages, or starting to emerge, in *Penium margaritaceum.* (C) Maximum likelihood phylogeny of GRAS proteins. The tree on the left includes GRAS proteins from five plant species: *P. margaritaceum* (green), *Marchantia polymorpha* and *Physcomitrella patens* (orange), and *Arabidopsis thaliana* and *Oryza sativa* (dark red). The collapsed clade (right) contains GRAS proteins of *P. margaritaceum* (green branches) and other algae (dark branches) based on transcriptome data. Pies indicate branches with bootstrap value < 90.

We looked for evidence of the morphological and physiological adaptations and key traits associated with terrestrialization through the reconstruction of gene family evolution, focusing on gene family expansions (Fig. 3A; Supplementary Table 5). A total of 11 expanded gene families (*P* < 0.05) in the ancestor of charophytes following separation from chlorophytes, were identified, while a particularly large number of expanded gene families was evident in the common ancestor of *P. margaritaceum* and land plants (N=124), compared to family expansions in earlier algal lineages. Given that charophytes contain paraphyletic lineages and that the Zygnematophyceae, including *P. margaritaceum*, is the closest lineage to land plants, this suggests stepwise gene family expansion (Catarino et al., 2016). The expanded gene families in *P. margaritaceum* were mostly associated with responses to stresses, such as water deprivation, cold, bacteria, and oxidative stress, as well as the production and signaling of the phytohormones abscisic acid (ABA), auxin (AUX), ethylene (ETH) and jasmonic acid (JA). Other substantially expanded gene categories related to protein phosphorylation and cell wall organization (Supplementary Table 6).

### Transcription factors and transcriptional regulators

The *P*. *margaritaceum* genome encodes 935 transcription factors (TFs) and 454 transcription regulators (TRs) (Supplementary Table 7 and 8), which is substantially more than those in either *K. nitens* (292 and 332, respectively) or *C. braunii* (496, 202). This contradicts the notion that morphological complexity correlates with the size of the TF/TR infrastructure (Lang et al., 2010) (Supplementary Fig. 4). The GRAS, NAC, LOB, bZIP, bHLH, WRKY and AP2/ERF-ERF families, all of which have been associated with abiotic stress responses in embryophytes, showed substantial expansions compared with other algal lineages (Fig. 3B). Moreover, the GRAS and BBR-BPC TF families, as well as specific subfamilies of the bHLH, WRKY, NAC and AP2/ERF-ERF families, may have originated in the Zygnematophyceae (Supplementary Fig. 5-7). For example, the *P. margaritaceum* genome encodes 15 proteins that are ancestral orthologs of the DREB subfamily of AP2/ERF-ERF TFs, but corresponding orthologs have not been found in other algae (Supplementary Fig. 7). In land plants, these regulatory proteins are involved in responses to abiotic stresses, such as cold, dehydration, salinity and heat (Agarwal et al., 2017).

Among the *P. margaritaceum* TF families, a notable feature is the remarkable large size of the GRAS family (291; Fig. 3B). Plant GRAS genes, named after *GIBBERELLIN-INSENSITIVE* (*GAI*), Repressor of *ga1-3* (*RGA*) and *SCARECROW* (*SCR*), together with SCR-LIKEs (SCLs), may have originated in bacteria (Zhang et al. 2012) and are present in land plants and some Zygnematophyceae (Engstrom, 2011; Delaux et al., 2015). In land plants, they have functionally diversified to regulate processes that are inherent to complex multicellular body plans and three-dimensional architecture, including meristem development, controlling cell division and expansion in roots and shoots, vascular development and seed maturation, as well as stress responses (Bolle, 2004; Ma et al., 2010). This functional divergence was hypothesized to occur after terrestrialization, as algal GRAS proteins form a monophyletic clade located outside of the land plant group (Hernandez-Garcia et al., 2019). However, we found that while most *P. margaritaceum* GRAS proteins clustered within the algal group (Fig. 3C), four clustered as an outgroup with a subgroup of land plant GRAS proteins (Fig. 3C). This suggests that GRAS proteins diverged in the Zygnematophyceae prior to the emergence of embryophytes.

### Phytohormone biosynthesis and signaling

In land plants, interlinked sets of phytohormone signaling pathways orchestrate the exquisitely complex cellular metabolic networks, developmental patterning, and systems that provide protection against environmental stresses, allowing exploitation of essentially all terrestrial habitats. The evolutionary origins of these phytohormones remains an intriguing question. A wide range of algal lineages, including charophytes, are capable of synthesizing and responding to a range of classical plant hormones but their physiological roles are typically not well understood (Ju et al., 2015; Lu and Xu, 2015; Ohtaka et al., 2017; Holzinger and Pichrtova, 2016). Moreover, the genomes of the two sequenced charophytes, *K. nitens* and *C. braunii*, do not encode the complete set of orthologs of land plant hormone biosynthesis and signaling pathways (Hori et al., 2014; Nishiyama et al., 2018) (Fig. 4; Supplementary Table 9; Supplementary Fig. 8-25). We investigated the genome of *P. margaritaceum* to find evidence of evolutionary innovations in the Zygnematophyceae associated with the F-box-mediated (auxin, JA, GA and strigolactone [SL]) and two-component (cytokinin [CK] and ETH) signaling pathways, as well as the ABA, salicylic acid (SA) and brassinosteroid (BR) pathways (Fig. 4).

**Figure 4.**
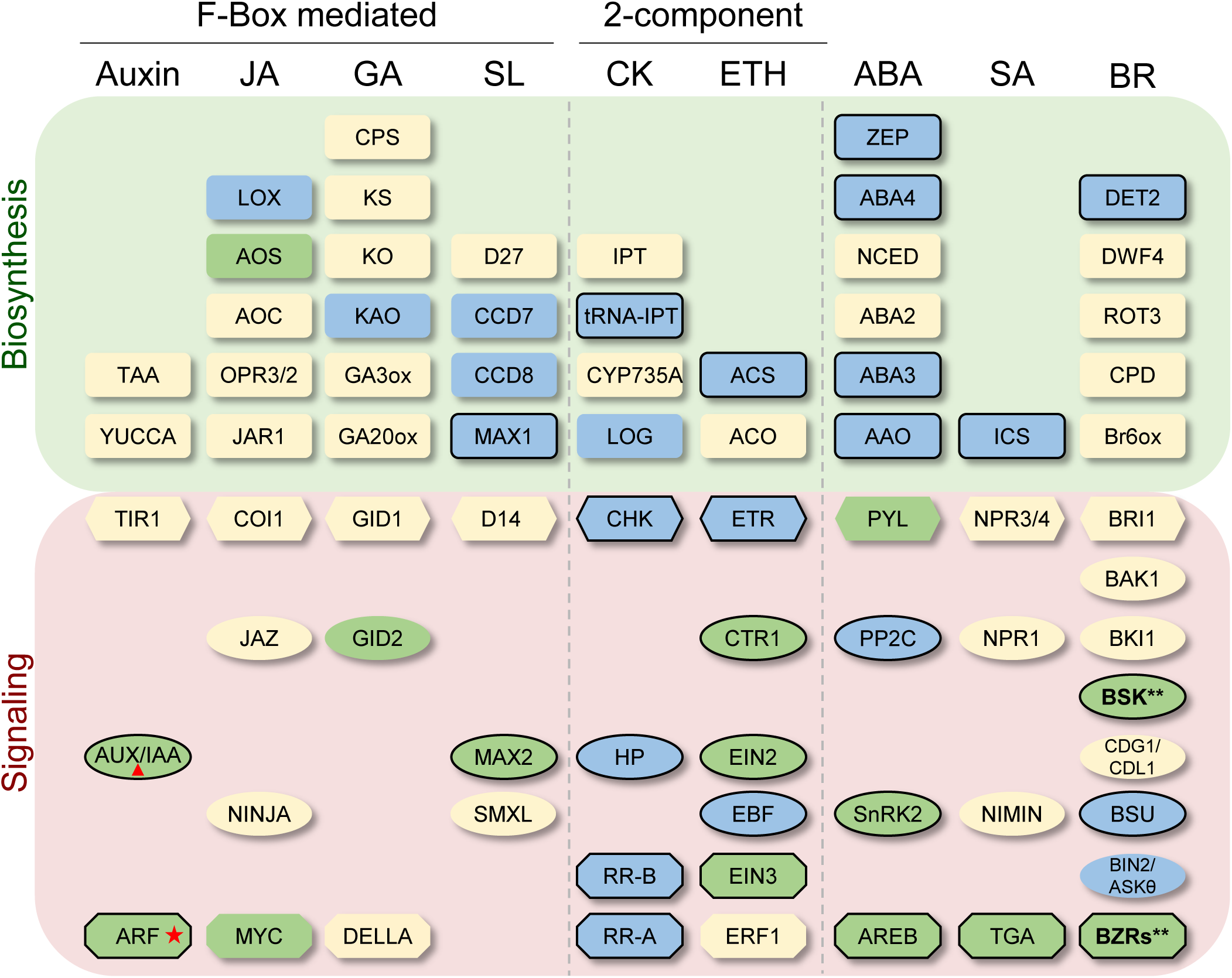
Genes involved in phytohormone biosynthesis and signaling. JA, jasmonic acid; GA, gibberellins; SL, strigolactone; CK, cytokinin; ETH, ethylene; ABA, abscisic acid (ABA); SA, salicylic acid; and BR, brassinosteroid. Rectangles, biosynthetic enzymes; hexagons, receptors; ovals, signal transduction components; and octagons, transcription factors. Shapes with a border indicate genes present in *Penium margaritaceum*. Double asterisks indicate genes that are present in *P. margaritaceum*, but not in earlier diverging lineages. The red triangle indicates a proto-domain I/II present in *P. margaritaceum* and the red star shows the proto A/B ARFs in *P. margaritaceum*. Blue shapes indicate genes already present in chlorophytes, and green and pale yellow shapes represent genes arising in charophytes and land plants, respectively.

Auxin coordinates a spectrum of growth and developmental processes via biosynthetic and signaling pathways that are conserved across land plants (Bowman et al., 2019). Various algal lineages also synthesize auxin and respond to its exogenous application, with cytological and structrual changes that are similar to those in land plants (Kiseleva et al., 2012; Ohtaka et al., 2017). However, both *P*. *margaritaceum* and *C*. *braunii* (Nishiyama et al. 2018) lack the primary auxin biosynthetic genes encoding tryptophan aminotransferase (TAA) and flavin-containing monooxygenases (YUCCA) (Supplementary Table 9). TAA and the paralogous family of alliinases are derived from a single land plant ancestor (Romani, 2017; Bowman et al., 2019) (Supplementary Fig. 8), and the YUCCA family is thought to have been acquired via horizontal gene transfer from bacteria to the ancestral land plant (Yue et al., 2012). We conclude that charophytes may use one of the alternative auxin biosynthetic pathways that have been proposed (Tivendale et al., 2014). In addition to the absence of a canonical auxin biosyntehtic pathway, neither *C. braunii* nor *P. margaritaceum* encode F-box genes that cluster with the land plant auxin receptor, *TIR1*, or its paralog *COI1*, which encodes a JA receptor, although *K. nitens* has one homolog that is likely the ancestor of both *TIR1* and *COI1* (Bowman et al., 2019) (Supplementary Fig. 9). Land plant auxin signaling involves binding of the TR AUX/IAA to the co-repressor TOPLESS, TIR1 and AUXIN REPSONSE FACTOR (ARF) TFs, through its I, II and PB1 domains, respectively (Leyser, 2018). *C. braunii* has two AUX/IAA genes and both lack domains I and II, while one of two *P. margaritaceum* AUX/IAA genes has prototypes of both domains (Supplementary Fig. 10A, Supplementary Table 9). This is reflected in a phylogenetic tree where the *C. braunii* AUX/IAA proteins cluster within a monoclade formed by non-canonical AUX/IAAs (NCIAAs), which lack domains I and II, whereas the *P. margaritaceum* homologs represent the ancestor of land plant canonical AUX/IAAs (Supplementary Fig. 10B). ARFs are ancient, predating the formation of a canonical auxin signaling network, and are categorized in land plants into three classes (A, B and C). The evolutionary history of these domains has not yet been fully resolved with the support of genome sequences (Martin-Arevalillo et al., 2019) (Supplementary Fig. 11A). The *C. braunii* genome encodes a single C-ARF, whereas the *P. margaritaceum* genome has two ARFs that cluster together as the ancestor of A/B-ARFs. Homology modeling revealed a high degree of protein structure conservation between the two *P. margaritaceum* ARFs and ARF1 (B-ARF) from the model land plant *Arabidopsis thaliana*, particularly in proximity to the dimerization domain (Supplementary Fig. 11B). This supports a model where both C and A/B classes were present in the common ancestor of *P. maragaritaceum* and land plants, and that loss of C-ARFs has occurred sporadically across charophyte lineages (Martin-Arevalillo et al., 2019).

Auxin transport and homeostasis rely on PIN and ABCB exporters, AUX1/LAX influx carriers and PIL proteins (Swarup and Bhosale, 2019). All of these transporters are found in the *P. margaritaceum* genome (Supplementary Table 9), while AUX1/LAXs and PILs are absent in *C. braunii* (Nishiyama et al., 2018). In land plants, polar auxin transport (PAT) is a key factor in the spatiotemporal control of development by asymmetric subcellular auxin distribution and is mediated by plasma membrane (PM) localized PIN proteins (Swarup and Bennett, 2014). The presence of PAT (Boot et al., 2012) and the polarized expression of PINs in charophytes (Żabka et al., 2016) suggest that PIN-mediated PAT may have originated in charophytes, although their relocalization to the plasma membrane from an ancestral form in the endoplasmic reticulum may have been key to the development of early land plants (Viaene et al, 2012). In conclusion, while it appears that the canonical auxin biosynthetic and signaling pathways were derived from the assembly and neofunctionalization of molecular interactions that existed in the ancestral land plant, the *P. margaritaceum* genome sequence has revealed additional core auxin signaling components that likely emerged in the Zygnematophyceae.

Genes required for the biosynthesis of JA and GA, and the associated canonical receptors and signaling elements, are absent from *P*. *margaritaceum* (Fig. 4; Supplementary Table 9), although both hormones have been detected in some charophytes (Kazmierczak and Rosiak, 2000; Hori et al., 2014). DELLA proteins are central repressors of GA-dependent processes and evolved from a subset of GRAS family proteins. However, while GRAS TFs are particualarly abundant in *P*. *margaritaceum*, none has the key N-terminal domain for interaction with the GID1 GA receptor (Hernández-García, et al. 2019). Our data are congruent with the idea that the DELLA proteins emerged in the land plant ancestor where they evolved a transciptional regualtory function, and were then recruited to form the GID1-DELLA signaling with the emergence of vascular land plants (Hernández-García, et al. 2019).

Similarly, there is evidence that canonical SL hormone signaling, which contributes to numerous developmental processes in land plants, including shoot branching, the initiation of lateral roots and leaf development, emerged in land plants through the recruitment of a pre-existing SL-based signaling system (Walker et al., 2019). We found that of the known biosynthetic pathway genes, *P*. *margaritaceum* only has an ortholog of *MAX1* (Fig. 4). Moreover, the *P*. *margaritaceum* genome encodes orthologs of only one SL signaling component, MAX2. While it has been reported that some charophytes have detectable levels of SLs and respond to their exogenous application (Delaux et al. 2015), the presence of SLs in charophytes has been questioned (Walker et al., 2019). The absence of a biosynthetic or signaling framework in *P. margaritaceum* is more consistent with the idea that SL synthesis orignated at the base of land plants.

In contrast to the F-box mediated hormones, there is considerable mechanistic conservation of the two-component hormone systems among streptophytes, consistent with early establishment deep in the lineage. There are structural differences between the cytokinin ligands synthesized by algae and angiosperms (Bowman et al., 2019), and the latter utilize adenylate-IPTs to generate *trans*-zeatin, while the class I tRNA-IPTs encoded by *P*. *margaritaceum* and other streptophytes produce *cis*-zeatin (Fig. 4; Supplementary Fig. 12). Notably, the *P*. *margaritaceum* genome lacks the LOG protein that, in land plants, converts inactive cytokinin nucleotides to biologically active forms (Kurakawa et al., 2007), while LOG is present in all of the other selected algal genomes, suggesting an alternative mechanism for cytokinin activation in *P*. *margaritaceum*. Key cytokinin signaling pathway components are found in *P*. *margaritaceum* and other algal genomes, except for the RR-A and RR-B response regulators, which are absent in *C*. *braunii*, again indicating functional substitution by other genes (Nishiyama et al., 2018). The ethylene pathway also has similarly highly conserved signaling elements, including ETR, CTR1, EBF, and EIN3, which are present in all three completed charophycean algal genome sequences (Fig. 4; Supplementary Table 9). The ancient evolution of ethylene as a signaling molecule was also demonstrated through physiological and transcriptome studies of *Spirogyra pratensis*, a filamentous close relative of *P. maragaritaceum* in the Zyngnematophyceae, showing regulation of abiotic stress responses, cell wall metabolism and photosynthesis (Ju et al., 2015; Van de Poel et al., 2016).

In land plants, ABA is associated with a range of developmental and physiological traits that are central to embryophyte life cycles, and with adaptive responses to the stresses and stimuli inherent in desiccating terrestrial habitats (Lievens et al., 2017; Eklund et al. 2018; Kollist et al., 2019; Kuromori et al., 2018). ABA can be synthesized in a diverse array of organisms via different biosynthetic routes (Siewers et al., 2006; Bowman et al., 2019). Two of the plant ABA biosynthetic genes, NCED and ABA2, are not found in *P. margaritaceum* and other selected algal lineages (Fig. 4; Supplementary Fig. 13 and 14). NCED is the rate-limiting enzyme (converting 9-*cis*-villa/neoxanthin to xanthoxin) and characterizes the plant-specific indirect ABA biosynthetic pathway (Hauser et al., 2011). The absence of these critical genes suggests that *P. margaritaceum* and other charophyte algae may employ a direct pathway, via farnesyl-diphosphate for ABA biosynthesis. This pathway has been identified in fungi (Siewers et al., 2006) and the associated genes are present in both algae and land plants (Supplementary Fig. 15). The land plant ABA signaling machinery involves several core components, including the ABA receptors PYR1/PYLs, negative regulators PP2C phosphatases and positive regulators SNRK2 kinases and AREB type bZIP TFs (Hauser et al., 2011). A homolog of PYR1/PYLs was not found in any of the algal genomes examined (Supplementary Fig. 16; Supplementary Table 9), nor in the transcriptomes of 15 Desmidiales genera, and was only present in two out of 13 genera of Zygnematales based on the transcriptome data (Ju et al., 2015; de Vries et al., 2018; Leebens-Mack et al., 2019), which is not congruent with the propsoal that PYL arose in the common ancestor of the Zygnematophyceae and land plants (de Vries et al., 2018). Nonetheless, other potential non-canoical ABA receptors in *A. thaliana*, such as ABAR and GCR (Cutler et al., 2010) were found in the algal genomes (Supplementary Fig. 17 and 18). Group A PP2C, group II and III SNRK2 and AREBs signaling components are all present in low copy numbers in *K. nitens* (Hori et al., 2014), *C. braunii* (Nishiyama et al., 2018), and *P. margaritaceum* (Supplementary Fig. 19-21; Supplementary Table 9), and an SNRK2 from *K. nitens* has been shown to transduce ABA-dependent signals when expressed in *A. thaliana* cells (Lind et al., 2015). This suggests an evolutionary retention by the first land plants of an ancestral ABA-mediated signaling and transcriptional regulatory module, which was then coupled via a novel receptor to land plant-specific ABA biosythetic machinery.

The only traces of a land plant-specific SA pathway in *P. margaritaceum* are an isochorismate synthase (ICS) homolog and TGA TFs; however, there is evidence of more extensive genetic innovation related to the BR phytohormone. The classical BR steroid hormone biosynthetic pathway includes DET2 and four members of the CYP85 clade of cytochrome P450 enzymes (Bak et al., 2011). DET2 orthologs are found widely in algae and land plants, but the four CYP85 enzymes are specific to vascular plants (Supplementary Fig. 22 and 23). BR regulates gene expression and plant development through a receptor kinase-mediated signal transduction pathway (Kim et al., 2009) and three out of the five kinases, including the BR receptor BRI1 and BAK1, are only present in land plants (Fig. 4; Supplementary Table 9). However, we found orthologs of BSK kinases, contrary to a recent report that BSKs were an embryophyte innovation (Li et al., 2019) (Supplementary Fig. 24), as well as BZR TFs (Supplementary Fig. 25) in *P. margaritaceum* but not in *K. nitens* or *C. braunii*. This suggests that these important components of the BR signaling circuitry, which governs cell elongation, interaction with other hormone networks, light signaling and stress responses in land plants (Sun et al., 2010; Ren et al., 2019), originated in the Zygnematophyceae.

### Cell walls and the diversification of extracellular structural polymers

The colonization of terrestrial habitats by embryophytes has been dependent upon the ability to synthesize complex cell walls that provide biomechanical support and protection against environmental stresses. Land plant primary walls are comprised of a core scaffolding of cellulose microfibrils embedded within matrices of interconnecting pectin and hemicellulose polysaccharides, together with glycoproteins (Burton et al., 2010; Popper et al., 2011; Dehors et al., 2019). However, immunological and biochemical studies suggest that the capacity to synthesize many of the polysaccharides of extant embryophyte walls evolved prior to the ancestral land plant, during divergence of the charophyte algae (Sørensen et al., 2011). Consistent with this idea, among the most remarkable examples of gene families showing expansion in *P. margaritaceum* are those encoding carbohydrate active enzymes (CAZymes; Cantarel et al., 2009) of the glycosyl hydrolase (GH), glycosyl transferase (GT), carbohydrate esterase (CE) and polysaccharide lyase (PL) classes, as well as carbohydrate binding modules (CBMs) and auxiliary activities (AAs) (Fig. 5A; Supplementary Table 10). CAZy enzymes are involved in diverse aspects of carbohydrate chemistry, including intracellular glycoconjugates, but notably include many that may be functionally associated with cell walls. There are relatively few, or in the case of PLs no, such genes in chlorophytes, and in every case there is a striking increase in abundance in *P. margaritaceum* compared with *C. braunii.* Moreover, in the cases of GTs and PLs there are more than in any of the green plant lineages. The large sizes of the classes typically reflect expansion within individual gene families (Fig. 5A).

**Figure 5.**
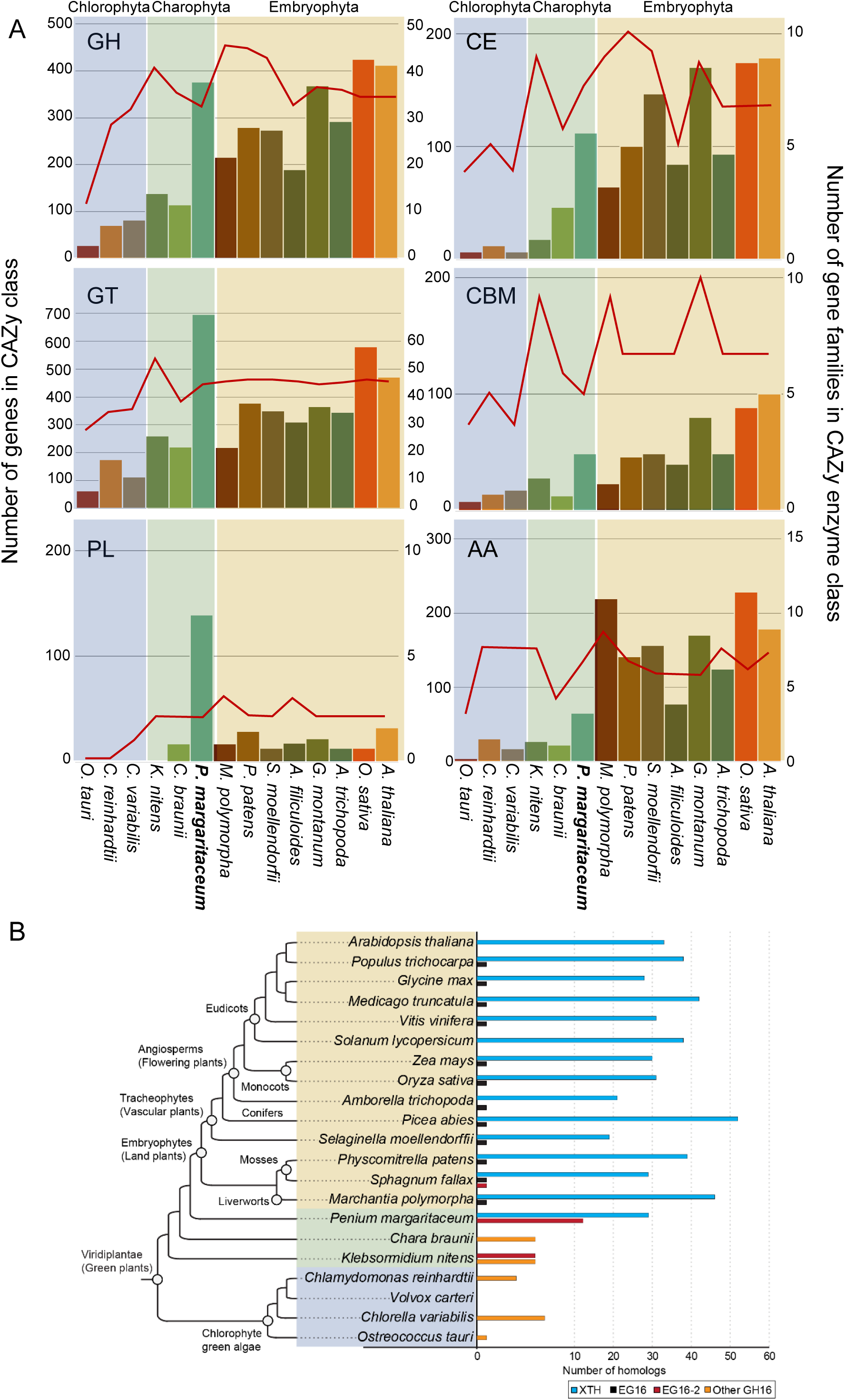
Carbohydrate active enzyme (CAZy) class composition. (A) Number of genes (left y axis; bars) and families (right y axis; lines) in the glycosyl hydrolase (GH), glycosyltransferase (GT), pectate lyase (PL), carbohydrate esterase (CE), carbohydrate binding module (CBM) and ancillary activity (AA) classes. (B) Census of EG16, EG16-2, XTH, and other GH16 homologs in the genomes of selected plants.

It might be expected that land plants with more complex body plans would have more extensive repertoires of CAZy proteins that orchestrate the restructuring of cell wall architecture during cell expansion and differentiation. However, *P. margaritaceum* has multiple families, associated with a range of cell wall polysaccharide substrates, which are considerably larger than those of *A. thaliana*. Particularly prominent examples of such gene family expansions are annotated as pectinases, such as the GH28 (polygalacturonase; 96 in *P. margaritaceum*, 67 in *A. thaliana*) and PL1 (pectate lyase; 139, 26) families, and GH16 (comprising xyloglucan tranglycosylase/hydrolase, XTH, and endo-glucanase 16, EG16; 41, 33) enzymes. The expansin family of cell wall loosening proteins (Cosgrove, 2015), which is not included in the CAZy grouping, show a similar trend (53, 35; Supplementary Table 11). Given that *P. margaritaceum* is unicellular, the particularly large size of these protein families is not explained by heterogeneity in wall architecture associated with different cell types or body plan complexity. Rather, it may reflect duplication and neofunctionalization resulting in differences in enzyme activities and properties, or in micro/nano-scale differences in spatial distribution.

Additionally, the high-level grouping of members in CAZy gene families can mask the emergence of novel enzymatic activities within distinct subgroups. Indeed, many GH and PL families are known to be “polyspecific”, encompassing several related, yet distinct, substrate specificities (Lombard et al., 2010; Viborg et al., 2019). GH16 is one such family, in which a unique subfamily of mixed-function plant endo-glucanases (comprising clades EG16 and EG16-2) has recently been delineated as a sister group to the XTHs (Elköf et al., 2013; McGregor et al., 2017; Behar et al., 2018). The presence of EG16-2 homologs in *P. margaritaceum* (12 genes) and in *K. nitens* (six genes; Fig. 5B; Supplementary Fig. 26), is concordant with the early evolution of this endo-glucanase subfamily (Behar et al., 2018), while the *P. margaritaceum* GH16 family composition suggests that XTHs originated in the Zygnematophyceae. The expansion of EG16 homologs in charophyte lineages is also striking because they are found exclusively as single genes in the later-diverging land plants (Fig. 5B; Behar et al., 2018). The complexity of GH16 family expansion and contraction is evident, but its functional significance will require elucidation by enzymology and structural biology data.

Another critical innovation for terrestrial plant life has been the elaboration of specific cell wall types with other classes of structural polymers to provide additional biophysical attributes for structural support and barrier properties. Examples include the phenylpropanoid polymer lignin in xylem vessel walls and the deposition of the structurally related lipid polyesters, cutin and suberin, in the hydrophobic cuticle of epidermal cells and the endodermis of roots, respectively (Fich et al., 2016; Renault et al., 2019). Algae do not have true cuticles, but a search of the *P. margaritaceum* genome for homologs of structural and regulatory genes that are known in *A. thaliana* to be associated with extracellular polyesters and wax cuticle components revealed traces of biosynthetic, transport and assembly frameworks (Supplementary Table 12). These encode enzymes involved in intracellular biosynthesis, as well as transporters and extracellular proteins that have been linked to extracellular lipid trafficking and cuticle assembly (Yeats and Rose, 2013). For example, homologs of cutin synthase (CUS) and BODYGUARD (BDG), which contribute to cuticle formation in land plants, are present in *P. margaritaceum* and *K. flaccidum.* However, other genes that are central to cuticle formation, such as that encoding glycerol-3-phosphate acyltransferase 6 (GPAT6), which forms monoacylglycerol cutin precursors, are unique to land plants, or present in far smaller numbers in algae. The quantitative and qualitative changes in cuticle-associated genes from chlorophytes to charophytes, and then again to land plants, (Supplementary Table 12), is consistent with the stepwise expansion and neofunctionalization of ancient core lipid biosynthetic machinery to synthesize structural lipid precursors, in conjunction with systems for their secretion. There is no evidence in *P. margaritaceum* or other charophytes of primordial cutin and suberin polyesters, and although wax-like lipid deposits have been reported in the cell walls of *K. nitens* (Kondo et al., 2016), the assembly of extracellular hydrophobic polymers was likely a land plant innovation.

There are parallels between the origins of cuticles and suberized walls, and the evolution of lignin, which is deposited in the secondary walls of specific tissues and cell types in land plants to provide structural reinforcement and protection against pathogens, and to limit water diffusion (Terrett and Dupree, 2019; Zhong et al., 2019). Lignin is synthesized through the phenylpropanoid pathway, and while lignin or lignin-like compounds have been reported in non-vascular plants, including charophytes, some of these likely resulted from misidentification of polyphenols and true lignin is specific to vascular plants (Weng and Chapple, 2010). The *P. margaritaceum* genome does not have genes that provide a canonical core phenylpropanoid pathway (Supplementary Table 13), including the enzyme phenylalanine ammonia lyase (PAL) at the entry point, and it has been suggested that it was acquired in land plants by horizontal gene transfer (Emiliani, et al., 2009). However, PAL is present in *K. nitens*, but not *C. braunii*, and other core phenylpropanoid biosynthetic genes show a similar ‘patchwork’ distribution among *K. nitens*, *C. braunii* and *P. margaritaceum* (de Vries et al., 2017; Supplementary Table 13). This suggests a complex evolutionary history in the production of soluble lignin-like compounds in charophytes, some of which are incorporated into the cell wall (Sørensen et al., 2011; Weng and Chapple, 2010). The development in charophytes and early land plants of mechanisms to secrete and assemble phenolic and aliphatic compounds likely gave rise to an increasingly diverse palette of protective extracellular biopolymers. These in turn paved the way for the formation of lignin, cutin, suberin and sporopollenin polymers that are found in the walls of extant land plants (Niklas et al., 2017; Renault et al., 2019).

Neo- and subfunctionalization of catalytically promiscuous enzymes, including those in the ancient shikimate pathway (Niklas et al., 2017), would provide metabolic plasticity, which is associated with the evolution and functional diversification of phenylpropanoid compounds. However, the absence in the genome of *P. margaritaceum* and other charophytes of clear candidates for key steps in the pathway leading to various phenylpropanoid compound classes suggests the existence of cryptic activities and novel enzymes. A notable example is flavonoids, which were originally thought to only exist in land plants, but have been identified in a few divergent algal lineages (Yonekura-Sakakibara et al., 2019). Among other functions, flavonoids provide protection against UV radiation, which would have been a major challenge for the first land plants, and so the evolutionary trajectory of flavonoid biosynthesis is of great interest. We definitively identified multiple classes of flavonoids in *P. margaritaceum* by mass spectrometry (Supplementary Fig. 27-30), consistent with the presence of biosynthetic routes that are commonly found in land plants (Supplementary Fig. 31). *P. margaritaceum* has a 4-coumarate:coA ligase (4CL) and, most notably, 11 homologs of chalcone synthase (CHS), which acts at the entry to flavonoid biosynthesis and is not present in earlier diverging plant lineages. Paradoxically though, *P. margaritaceum* has neither PAL nor a cinnamate 4-hydroxylase (C4H) in the same cytochrome P450 subfamily (CYP73A) as the C4H genes of land plants (Yonekura-Sakakibara et al., 2019) (Supplementary Table 13). Thus, there is no clear mechanism to synthesize cinnamic acid and coumaric acid, which are intermediates in the formation of the coumaroyl-CoA substrate for CHS (Supplementary Fig. 31). Some of the genes functioning downstream of CHS, such as chalcone isomerase (CHI) and flavanone 3-hydroxylase (F3H), which lead to the spectrum of flavonoid compounds, are also absent. Whether these apparently missing steps are catalyzed by proteins in the same superfamilies as those of extant land plants, but are more distantly related, or they represent alternative biosynthetic routes to the same product, remains an open question.

### Effects of terrestrial abiotic stresses on cellular responses and transcriptome dynamics

To gain further insights into the molecular processes and adaptations that allow *P. margaritaceum* to tolerate the severe physiological challenges imposed by its ephemeral habitat, we conducted transcriptome profiling of responses to two archetypal terrestrial environmental factors associated with a terrestrial habitat: desiccation (DE) and high light (HL), as well as a combination of the two (HLDE). HL had no notable effect on cellular or chloroplast morphology but DE, imposed by placing the cells on cellulose sheets, and to a lesser degree HLDE, induced asymmetric cell elongation and disruption of the characteristic lobed chloroplast architecture. (Fig. 6A). DE and HLDE treatments also caused substantial accumulation of starch in the chloroplast, as well as the formation of large cytoplasmic vacuoles, some of which showed evidence of autophagy (Supplementary Fig. 32). Starch degradation and biosynthesis have both been observed as abiotic stress responses in different plant taxa (Thalmann and Santelia, 2017), and in land plants, autophagy is associated with stress tolerance, the recycling of organelles and macromolecules, and ROS scavenging (Signorelli et al., 2019).

**Figure 6.**
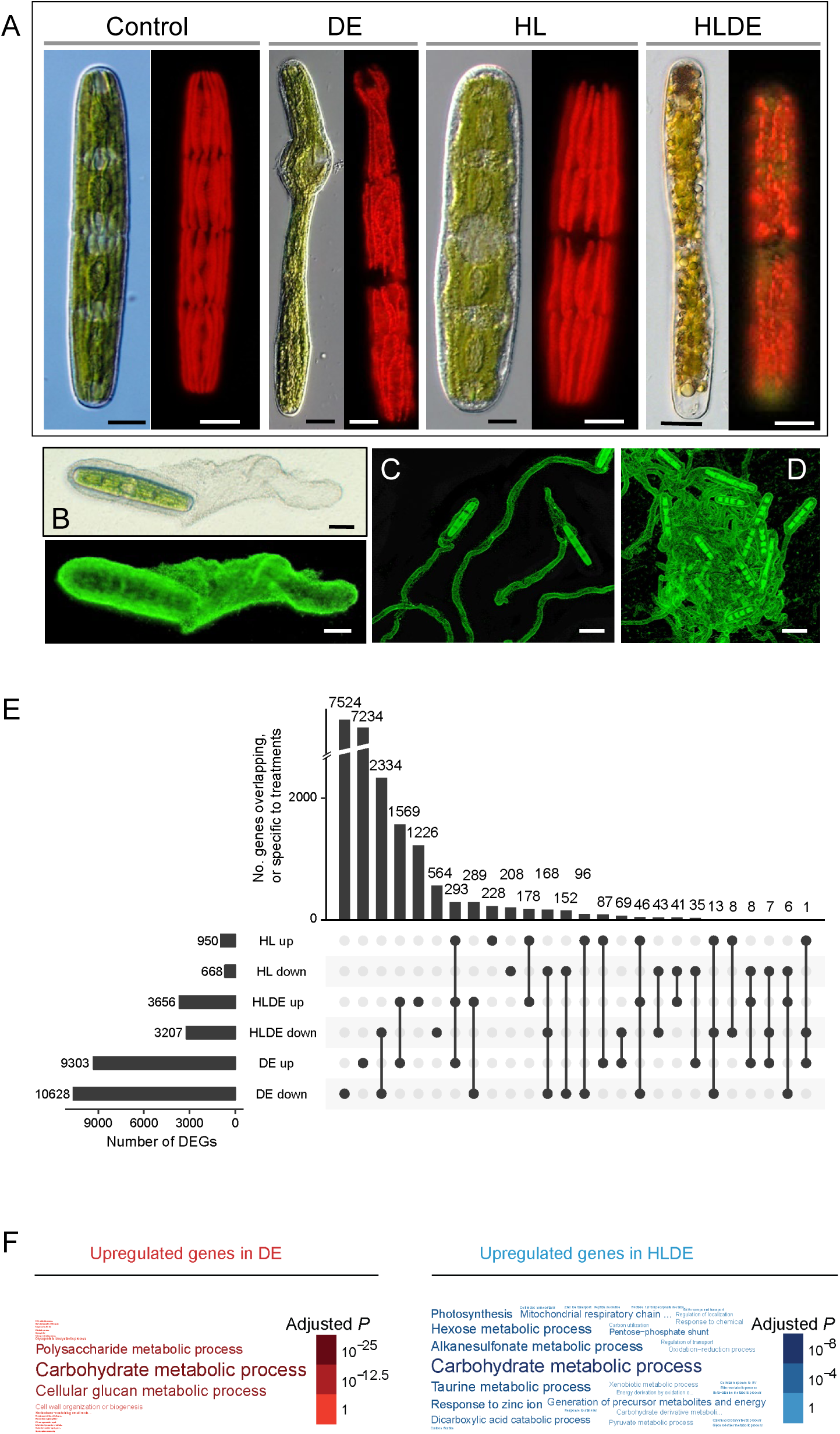
Responses of *Penium margaritaceum* to high light and desiccation stress. (A) Morphology of *P. margaritaceum* cells grown under control, high light (HL), or desiccating (DE) conditions, or a combined treatment (HLDE), imaged using differential interference contrast microscopy (DIC; left panel of each condition) and fluorescent light microscopy (FLM; right panels). (B) DIC (top) and confocal laser scanning microscopy (CLSM) image (bottom) after labeling with Fluoresbrite beads, showing mucilage secretion predominating from one pole of the cell, resulting in propulsion. (C) Gliding trails in control cells. (D) Dense cellular aggregation following HL treatment. (E) Gene expression analysis, based on RNA-seq data, of HL, DE, and HLDE treated *P. margaritaceum* cells compared with cells grown under control conditions. (F) Word clouds of gene ontology (GO) terms in the ‘biological process’ category related to differential gene expression under DE and HLDE conditions.

A major structural and behavioral effect of all three treatments was the production of large quantities of mucilage (Fig. 6B). Many zygnematophycean algae secrete large amounts of extracellular polysaccharide mucilage through their cell walls, creating an extensive hydroscopic sheath. This material, also referred to as extracellular polymeric substance (EPS), has many functions that would provide an evolutionary advantage in semi-terrestrial habitats: anti-desiccation; a matrix for conjugation; a biofilm for communication with other microorganisms; and a propulsion mechanism where secretion from one pole of the cell allows directional gliding motility (Boney, 1981; Brook, 1981; Fisher et al., 1998; Oertel et al., 2004; Domozych et al., 2005; Kiemle et al., 2007; Domozych and Domozych, 2008). Mucilage production, which results in cell gliding behaviors (Supplementary Video 1 and 2) increased substantially within a few minutes of applying the HL, DE and HLDE treatments (Fig. 6C,D; Supplementary Fig. 33). Under DE and HLDE conditions, the EPS trails were more densely packed, leading to cell aggregation. This may be beneficial to an ephemeral alga whose short active growth period in the summer is defined by the correlation of high light with drying conditions in shallow wetlands. The tight packing of EPS trails and cells would provide a means of enhancing water retention in a hydroscopic mass under drying conditions.

Consistent with the degree of the morphological and cytological changes, transcriptome profiling of *P. margaritaceum* revealed a greater response to DE than to the other two treatments (Fig. 6E; Supplementary Fig. 34), with 9,303 and 10,628 genes up- and down-regulated, respectively. Most of the DE-related differentially expressed genes (DEGs; 78% of the up-regulated and 71% of the down-regulated) were DE-specific. HL had the least impact on transcript profiles, while the combined treatment had an intermediate effect. Under HLDE, 51% and 78% of the up- and down-regulated genes, respectively, showed the same expression patterns as under the DE treatment. These results suggest that elevated light levels alleviate the impact of DE stress.

Gene ontology (GO) enrichment analysis of the DEGs (Supplementary Table 14-19; Supplementary Fig. 35) showed that the predominant categories of genes up-regulated by all three treatments are related to carbohydrate metabolism. The HL treatment caused an induction of genes related to central carbon metabolism, while photosynthesis related pathways and associated chloroplast related genes were significantly down-regulated. This is consistent with suppression of photosynthesis to prevent cellular damage caused by HL-induced ROS, as occurs in land plants (Rossel et al., 2007). Complex networks of *P. margaritaceum* genes were identified as being regulated by DE or HL, including representatives of families that are not present in other sequenced algal genomes. A notable example was GRAS, which corresponded to the TF family with the greatest number of DEGs under DE (50 and 66 induced and repressed, respectively), while none was differentially expressed under HL, consistent with an ancestral role in abiotic stress responses. In addition, 12 of the 15 *P. margaritaceum* DREB TFs (Supplementary Fig. 7), which are also not found in other algal linages, were responsive to DE (three up-regulated and nine down-regulated), consistent with a role in adaptations to increasingly terrestrial habitats

One of the most prominent transcriptome responses was a major up-regulation by DE of genes annotated as being involved in polysaccharide metabolism and cell wall biosynthesis (Fig. 6F; Supplementary Table 14 and 18). These include members of various GT classes, glycan synthases, and transglycosylases that function in the synthesis of diverse land plant cell wall polymers, including cellulose, xylan and pectins. It might be expected that the transcriptome profiles reflect the biosynthesis of the mucilage that was induced in large quantities. An analysis of the polysaccharides in the mucilage secreted following HL or DE treatments (Supplementary Table 20 and 21) revealed that they are quite distinct from those of the *P. margaritaceum* cell wall (Sørensen et al., 2011), as well as showing compositional differences in response to different treatments, and so these gene sets may provide useful targets for future studies of the biosynthesis and function of the mucilage polymers.

The transcriptome profiles also suggested that substantial cell wall remodeling occurred in response to DE. Large proportions of several of the families associated with cell wall loosening and degradation (48%, 77%, 68% and 58% of GH28, PL, GH16, and expansin genes, respectively) were up-regulated in DE stressed cells. Notably, only 1-3% and 6-23% of genes in these families were up-regulated under HL or HLDE treatment, again suggesting that the effects of DE were offset by higher light conditions. Congruent with the upregulation of GH28 and PL pectinase genes, immunological analysis with a monoclonal antibody (JIM5) that recognizes the pectin polymer homogalacturonan (HG), showed that the application of DE or HLDE stress caused major changes in the pectin architecture at the site of wall expansion at the isthmus zone (Supplementary Fig. 36A). This was confirmed by ultrastructural observations whereby the HG lattice was significantly reduced, leaving the inner cellulosic wall layer (Supplementary Fig. 36B,C). This alteration most likely compromises the structural integrity of the wall, resulting in the unusual shapes of cells grown under these stress conditions. These results add to growing evidence that abiotic stresses, such as desiccation, cause remodeling of the cell wall in both charophytes (Herburger and Holzinger, 2015; Holzinger and Pichrtova, 2016) and land plants (Tenhaken, 2015). The major expansion of cell wall modifying protein families in *P. margaritaceum*, together with their upregulation and turnover of their substrates in response to desiccation, highlights the importance of a dynamic primary cell wall to withstand changing osmotic conditions, and the significance of habitats such as transient wetlands in land plant evolution.

## DISCUSSION

Approximately 500 Mya, an ancestor of the modern day Zygnematophyceae emerged from a transient freshwater habitat and colonized a barren terrestrial surface. Subsequent evolutionary “tuning” gave rise to the great diversity of land plants that has ultimately transformed the natural history and biogeochemistry of the planet. The *P. margaritaceum* genome sequence confirms the Zygnematophyceae as the sister lineage of land plants, and has the hallmarks of a dynamic source of genetic innovation, with abundant TEs and the emergence, or major expansion, of gene families and regulatory systems that are associated with terrestrialization. These include a large compendium of regulatory TF families and components of phytohormone signaling networks that govern stress responses and cell morphology in embryophytes.

The genome also provides evidence that several key land plant characteristics found in *P. margaritaceum* may have been critical pre-adaptations for the successful transition to life in a terrestrial habitat. Key among these are an extensive machinery for the synthesis, secretion and remodeling of the polysaccharide cell wall, much of which originated prior to the first true land plant. This is exemplified by the substantially expanded repertoire of genes involved in the metabolism of pectins. These apparently ancient macromolecules contribute to cell expansion and cell differentiation, as well as forming the middle lamella that mediates intercellular adhesions, allowing tissue and organ formation in land plants (Zamil and Geitmann, 2017; Cosgrove, 2014). Addionally, while most terrestrial plant life requires more extensive depsoition of hydrophobic biopolymers to reinforce the walls of specialized cells, the origins of their building blocks can increasingly be traced back to aquatic ancestors.

The unicellular habit of *P. margaritaceum* repesents a major evolutionary reduction that affords significant advantages to life in aquatic habitats that experience periodic drying. Small size, rapid cell division, simple conjugation-based sexual reproduction and the ability to withstand desiccation-based stress, and the synthetic machinery to secrete large amounts of water-retaining mucilage, provide a more efficient means to survive in shallow wetlands than the multicellular habits and complex reproductive strategies displayed by other late-divergent charophytes. Ancient zygnematophyceaen algae living in isolated freshwater pools were well adapted to make the move to life on land. It is important to note that all zygnematophyceaen taxa are believed to be derived from multicellular ancestors (Delwiche and Cooper, 2015). Upon initial land colonization, a reversion to a multicellular form that would provide a greater surface area for photsynthesis and the absorption of minerals and water from the “new” substrate most likely occurred.

More elaborate plant body plans evolved independently in different lineages of the Streptophyta, with members of the Charophyceae taking advantage of the buoyancy provided by their exclusively aquatic environment and the Coleochaetophyceae using a highly compact thallus and unique sensory hairs to live in semi-aquatic and terrestrial habitats. However, it was the Zygnematophyceae that evolved significant thallus reduction, fast growth rates and simplified conjugation-based sexual reproduction, to thrive in transient freshwater wetlands that most likely dominated Earth’s land surfaces over 500 Mya. As important, these and other characteristics described in this study were critical to successfully colonizing land. Once established, proliferation and subsequent evolutionary events led to the land plants.

## MATERIALS AND METHODS

### General culture conditions

*Penium margaritaceum* Brébisson (Skidmore College Algal Culture Collection) was maintained in sterile 100 mL liquid cultures of Woods Hole medium (Nichols, 1973) supplemented with soil extract (WH soil or WHS: soil extract obtained from Carolina Biological, USA), pH 7.2 at 18 ± 2°C in a photoperiod of 16 h light/8 h dark with 74 μmol photons m^−2^ s^−1^ Photosynthetic Photon Flux of cool white fluorescent light. Subcultures were made every 10 days and 10-14 day old cultures were used for all experiments.

### Stress conditions

Stress cultures were maintained in 50 mL aliquots of WHS in sterile 150 × 15 mm plastic Petri dishes. Cultures (5 mL; 2,000 cells mL^−1^) were added to each dish, which was then sealed with surgical tape and placed under high light (HL; 150 μmol photons m^−2^ s^−1^, 18 +/−2°C), or control (74 μmol photons m^−2^ s^−1^, 18 +/−2°C) conditions. For the desiccation (DE) stress experiments, 150 × 15 mm plastic Petri dishes were filled with 50 mL of WHS containing 2% agarose (Sigma. A-1296) and allowed to cool. Two sterile 80 mm diameter cellulose sheets (325p; AA Packaging Limited) were added to each plate and 1 mL of concentrated cell culture (∼5,000 cells mL^−1^) was spread onto the sheets. The plates were then sealed and placed under control conditions or HL to produce high light plus desiccation (HLDE) conditions. Cells from three independent treatments or the control were collected after 14 days by centrifugation at 1,500 × *g* for 1 min, the pellets washed three times by resuspension in sterile WHS, shaking and centrifugation, and then frozen in liquid nitrogen. For DE experiments, the cells were scraped off the cellulose sheets and frozen in liquid nitrogen.

### Cell labeling and imaging

Cells grown in liquid culture under control or stress conditions were washed three times with WHS, centrifuged (400 × *g* for 1 min) and the cell pellet was gently resuspended in WHS (1,000 cells mL^−1^ +/− 50). Samples (200 µL) were placed in the center of a single-welled polytetrafluoroethylene printed slide (Electron Microscopy Sciences, Hatfield, PA, USA) and mixed with 50 µL of a 100 µg/mL solution of Fluoresbrite Plain YG 0.5 µm Microspheres, or with 0.5 µm Polybead Polystyrene Microspheres (Polysciences, Warrington, PA, USA). The mixed suspensions were covered with a 22 × 22 mm glass coverslip and imaged with either light microscopy (LM; Olympus BX-60 or IX-83 equipped with both wide field fluorescence and differential interference contrast optics), or confocal laser scanning microscopy (CLSM; Olympus Fluoview 1200 CLSM). Single images or Free Run time lapse video clips were acquired with Olympus DP-73 cameras. For some experiments, the slides were placed in a moisture chamber comprising a glass Petri dish with a layer of wet filter paper. The chambers were placed under the control and HL conditions for various periods of time and then viewed with LM or CLSM. To image the mucilage in DE cultures, cell aggregates were removed from the surface of desiccation cultures, placed on the slides as above and a drop of 0.5 mg/mL Fluoresbrite bead solution was placed on the cell aggregate. The coverslip was then added and the slide viewed with LM or CLSM. To visualize starch, cell pellets were resuspended in growth medium and stained for 5 min with 1% v/v iodine then washed before imaging. Immunolabeling of cell wall pectin followed the protocol of Rydahl et al. (2015) with the anti-HG monoclonal antibody, JIM5 (Knox et al., 1990). Immunolabeling of the mucilage (EPS) (Fig. 1D) was as described in Domozych et al. (2005).

For transmission electron micrograph (TEM) imaging, cell suspensions were spray frozen into liquid propane cooled with liquid nitrogen. Freeze substitution and embedding followed the protocol of Domozych et al. (2007) and 80 nm sections were cut on an ultramicrotome (Leica), stained with conventional uranyl acetate and lead citrate and viewed with a Zeiss Libra 120 transmission electron microscope. For scanning electron microscopy imaging (SEM), cells were collected by centrifugation, placed on nitrocellulose paper attached to a JEOL Cryostub (JEOL, USA) and frozen in liquid nitrogen then imaged with a JEOL 6480 LV SEM under low vacuum conditions (10 kv, spot size 30).

### DNA and RNA extraction

Cells were grown for 14 days in sterile 125 mL flasks containing 75 mL of WHS under the conditions described above. Cells were then collected by centrifugation, and used as a source of RNA (Wan and Wilkins, 1994). The quality was confirmed using an Agilent BioAnalyzer (Agilent, Santa Clara, CA, USA). RNA for ISO-seq was extracted from cells grown for 3 days in mating inducing media (Sørensen et al., 2014) using the RNeasy Mini kit (Qiagen, USA). Nuclei were isolated from the cell pellets (Raimundo et al., 2018) and used as a source of genomic DNA as this yielded far better quality preparations. DNA quality was verified using a Nanodrop™ 2000 Spectrophotometer (Thermo Fisher Scientific, USA) and an Agilent BioAnalyzer (Agilent, Santa Clara, CA, USA).

### Library construction and sequencing

Five paired-end libraries were constructed for genome sequencing, of which three were prepared with the Illumina TruSeq DNA PCR-Free Prep kit, one with the Illumina Genomic DNA Sample Prep kit and one with the Kapa Hyper kit (Kapa Biosystem, Roche). Three mate-pair libraries were constructed using Illumina Nextera Mate Pair Library Prep kit with insert sizes ranging between 2-4 kb, 5-7 kb and 8-10 kb, respectively (Supplementary Table 1). All libraries were sequenced on an Illumina HiSeq 2500 system in paired-end mode.

Strand-specific RNA-Seq libraries were constructed for each sample as previously described (Zhong et al., 2011) and sequenced on an Illumina HiSeq 2500 system in paired-end mode. A non-size-selected SMRTbell (Pacific Biosciences, USA) library from the total RNA was constructed using the manufacturer’s Iso-Seq protocol and sequenced in two SMRT cells on the PacBio Sequel platform (v2.0 chemistry).

### Sequence processing, *de novo* assembly and quality evaluation

Genomic paired-end reads were processed to remove adaptors and low-quality bases using Trimmomatic (Bolger et al., 2014) (version 0.32) with parameters “TruSeq3-PE-2.fa:2:30:10:1:TRUE SLIDINGWINDOW:4:20 LEADING:3 TRAILING:3 MINLEN:40”. Mate-pair reads were cleaned with the ShortRead package (Morgan et al., 2009) to remove the junction adaptor sequences formed during library construction and the trailing bases.

To assemble the genome, we first searched for optimal k-mer size. Since the *P. margaritaceum* genome is highly repetitive (Supplementary Fig. 1A), large k-mer size can help resolve repetitive regions with a trade-off of increasing the number of unique k-mers, which requires more computational resources. For *P. margaritaceum* genome assembly, memory efficiency was a major bottleneck for most of the popular assemblers attempted, including MaSuRCA (Zimin et al., 2013), SPAdes (Bankevich et al., 2012), ALLPATHS-LG (Gnerre et al. 2011), ABYSS 2.0 (Jackman et al., 2017) and w2rap-contigger (Clavijo et al. 2017). All of them failed after running days to weeks on a 1Tb memory machine and some even failed on a 3Tb memory cluster. The final genome assembly was generated by SOAPdenovo2 (Luo et al., 2012) (version 2.04) on a 1Tb memory machine with kmer size set to 127. The redundant contigs were removed using BLASTn (coverage ≥90% and identity ≥99%) and the remaining contigs were assembled into a scaffold using the built-in module of SOAPdenovo2, with reads from all the mate-pair and paired-end libraries (parameters ‘-F -u’). Gaps in the resulting scaffolds were filled using GapFiller v1-10 (Nadalin et al. 2012) with all paired-end reads.

Additional steps were taken to improve the contiguity and gene space of the assembly. First, the genome was also assembled successfully by MEGAHIT v1.1.1-2-g02102e1 (Li et al., 2015) using discrete k-mer sizes (121, 139, 159, 179, 199, 219, 239 and 249). Although the overall quality of the MEGAHIT assembly (contig N50: 498bp; scaffold N50: 696 bp) was not better, some long contigs were used for further scaffolding or gap filling of the SOAPdenovo2 assembly. MEGAHIT contigs > 2 kb were used in a BLAST search against the SOAPdenovo2 assembly (identity > 99%; minimal HSP length > 100 bp). Gapped scaffolds uniquely spanned by MEGAHIT sequences were filled, or replaced if they were contained by MEGAHIT sequences. When one MEGAHIT contig was aligned with two SOAPdenovo2 scaffolds, and the two alignments in the MEGAHIT contig did not overlap, the SOAPdenovo2 scaffolds were joined with gaps (100 Ns) if: the anchors 1) were located on each contig at the termini; and 2) were ≥100 nucleotides. Second, the genome assembly was further polished with assistance of transcriptome *de novo* assembly. We aligned the transcripts to the genome assembly using GMAP (Wu and Watanabe 2005) (version 2017-10-12) with parameters “--min-identity=0.95 --max-intronlength-middle 50000 -L 100000”. If a transcript was uniquely mapped onto two scaffolds and the alignments met the following criteria, the two scaffolds were joined with gaps (100 Ns): a) individual alignment length > 100 bp; b) total alignments should cover > 80% of the transcript; and c) alignments on the scaffolds should be located within 5 kb of the closest terminus. Consequently, 3,971 scaffolds (∼111 Mb) were successfully connected. Lastly, transcripts not covered by the SOAPdenovo2 assembly but with solid homologs in the NCBI protein database were mapped to the MEGAHIT assembly, and the aligned MEGAHIT contigs, if not redundant, were added to the SOAPdenovo2 assembly.

To correct base errors in the assembly, the variants were called with all paired-end reads. Briefly, cleaned reads were aligned to the assembly using BWA 0.7.15-r1140 (Li and Durbin 2009) and valid alignments (mapping quality ≥ 40; properly paired) were used for SNP calling by FreeBayes (v1.2.0) (Garrison and Marth 2012) with parameters “-q 0 -F 0.2 -p 1”. The variants were filtered by BCFtools (Narasimhan et al., 2016) using the stringent criteria “QUAL>30 & TYPE=‘snp’ & AO/DP >0.5 & DP>=30 & DP<=300 & MQM>30 & SAF>1 & SAR>1 & RPR>1 & RPL>1”. The resulting high-confidence variants were used for base correction of the assembly. The assembly was further checked for redundancy as described above. Scaffolds with BLASTn hits (E-value < 1e-5) from non-eukaryotic organisms were manually examined to exclude contamination.

To assess the quality of the assembled genome, Illumina genomic and RNA-Seq reads were mapped to the assembly through BWA or HISAT2 (Kim et al., 2015) (v2.1), respectively. K-mer based analysis were carried out with KAT (Mapleson et al., 2016) (V2.4.1).

### PacBio Iso-Seq and Illumina RNA-Seq data analysis and transcriptome *de novo* assembly

Long reads produced by the PacBio Sequel platform were processed using modules in the SMRTLink package (v5.1; PacBio), to generate full-length refined consensus isoforms. Circular consensus sequences (CCSs) were obtained from the ‘ccs’ module using the parameters “-- minPredictedAccuracy=0.75, MinFullPasses =0 and --minLength=100”. CCSs containing poly(A) signal, 5’ and 3’ adapters were then identified, and the adapters and poly(A) sequences were trimmed to create full-length non-chimeric reads (FLNC). The retained FLNC reads were iteratively classified into clusters to build the consensus sequences, which were then polished by Quiver (Chin et al., 2013) with the minimum accuracy rate set to 0.99. Base errors in the polished isoforms were further corrected by Illumina RNA-Seq reads using LoRDEC v0.8 (Salmela and Rivals, 2014) with kmer length set to 19. The LoRDEC-corrected isoforms were used to reconstruct the coding regions of the *P. margaritaceum* genome using Cogent v3.5 (https://github.com/Magdoll/Cogent). To build gene clusters, all the isoforms were aligned to the *P. margaritaceum* coding genome (the collection of coding sequences generated by Cogent) by minimap2 with parameters “-ax splice -uf” (Li, 2018), and the resulting alignments were then processed by cDNA_Cupcake (https://github.com/Magdoll/cDNA_Cupcake) to collapse isoforms.

Illumina RNA-Seq reads were processed with Trimmomatic (Bolger et al., 2014) to remove adaptor and low-quality sequences. Reads aligned to the ribosomal RNA database (Quast et al., 2013) were discarded. The remaining cleaned reads were subjected to Trinity (Grabherr et al., 2011) for *de novo* assembly with the minimum kmer coverage set to two. To remove redundancy, Trinity assembled contigs were further clustered by iAssembler (Zheng et al., 2011) with a sequence identity cutoff of 97%.

### Repeat annotation

A species-specific repeat library was built following the advanced repeat library construction tutorial described in Campbell et al. (2014). Specifically, LTRharvest (Ellinghaus et al., 2008) (v1.5.9; parameters ‘-minlenltr 100 -maxlenltr 6000 -mindistltr 1500 -maxdistltr 25000 -mintsd 5 -maxtsd 5 -motif tgca -vic 10’) and LTRdigest (http://genometools.org/tools/gt_ltrdigest.html) were used to identify long terminal repeat (LTR) retrotransposons, and MITE-Hunter (Han and Wessler, 2010) (v11-2011; parameters ‘-n 30’) to identify miniature inverted transposable elements (MITEs) in the genome assembly. The identified LTRs and MITEs were used to mask the *P. margaritaceum* genome using RepeatMasker (v4.0.7; www.repeatmasker.org), and the unmasked genomic sequences were analyzed by RepeatModeler (v1.0.11; http://www.repeatmasker.org/RepeatModeler.html) to identify novel transposable elements (TEs). All identified repeat sequences were searched against the Swiss-Prot database (www.uniprot.org/) using BLASTx with an E-value cutoff of 1e-10, and repeats matching non-TE proteins in the database were excluded. To annotate the repeats, we used a modified approach similar to that implemented in the RepeatClassifier module of RepeatModeler. First, we ran the RepeatClassifier on all identified repeats to get a summary statistic of BLASTx matches against a TE protein dataset provided by RepeatMasker. Repeats were categorized based on the classification of best hits (filtered by E-value < 1e-3 and alignment length > 150 nucleotides or 50 residues). The unclassified repeats were scanned by RepeatMasker with all eukaryotic TEs from the Repbase database (version 20170127). Best hits, with alignments > 200 nucleotides, were kept for TE family assignment. Simple repeat annotation was performed with an independent run of the TRF program (Benson, 1999) (v 4.09; parameters: “2 7 7 80 10 50 2000 -h -f -d -m 1 -l 10”).

### Estimation of LTR retrotransposon insertion time

The long terminal repeat (LTR) sequences of each identified full-length retrotransposon were aligned with MAFFT (Katoh and Standley, 2013) (v7.313; parameters: “--maxiterate 1000 – localpair”), and the genetic distance was estimated using the distmat program from the EMBOSS package (Rice et al., 2000) with the Kimura method. The insertion time (T) of the LTR retrotransposons was calculated according to the formula T=K/2r, where K is the genetic distance and r is the nucleotide substitution rate, which was estimated to be 7.0 × 10^−9^ substitutions per site per year in *A. thaliana* (Ossowski et al., 2010).

### Gene prediction

We predicted protein-coding genes in the *P. margaritaceum* genome using the MAKER-P pipeline (Cantarel et al., 2008) (v2.31.10), which integrates gene models derived from three sources of predictions: *ab initio* prediction, protein homology based evidence, and transcript evidence. Three *ab initio* predictors, GeneMark-ES v3.51 (Lomsadze et al., 2005), SNAP v2006-07-28 (Korf, 2004) and AUGUSTUS v3.3 (Stanke et al., 2006), were incorporated in the MAKER-P pipeline. Proteomes of 18 plant species across the Viridiplantae were used to identify protein homology. To prepare the transcript evidence, RNA-Seq reads were assembled using Trinity v2.6.6 (Grabherr et al., 2011) under both *de novo* and genome-guided modes. The resulting two assemblies, as well as an additional source of transcript structures obtained from StringTie 1.3.3b (Pertea et al., 2015), which used RNA-Seq alignments to the *P. margaritaceum* genome by HISAT2 (Kim et al., 2015) (v2.1), were supplied to the PASA pipeline v2.2.0 (Haas et al., 2003) to build a comprehensive transcriptome assembly. Protein-coding regions of the PASA assembly were predicted by TransDecoder (https://github.com/TransDecoder/TransDecoder/wiki) and confirmed through homology search against the Pfam (Bateman et al., 2004) and Uniref (Suzek et al., 2014) protein databases, PASA predicted gene structures were used as a training set for AUGUSTUS and also served as an independent prediction considered by the MAKER-P pipeline. The Trinity assemblies, combined with 29,220 expressed sequence tags (ESTs) from the NCBI nucleotide database, were fed to the MAKER-P pipeline as transcript evidence.

The final MAKER-P gene models were compared to the Pfam database to exclude those containing TE-related domains. Genes with expression value RPKM (reads per kilobase of exon model per million mapped reads) ≥ 1 in combined RNA-Seq data or having at least one valid hit in any knowledge databases (nr/Pfam/InterPro/PANTHER) were classified as high-confidence genes, whereas the rest of the predicted genes were grouped into a low-confidence gene set.

### Development of the master gene set

To further improve gene completeness, a master gene set was created by incorporating PacBio Iso-Seq full-length transcript isoforms and Trinity assembled transcripts into the high-confidence (HC) gene set. Transcript sequences were subjected to SeqClean (https://sourceforge.net/projects/seqclean) for polyA tail trimming and rRNA sequence removal. Coding regions of the transcripts were identified by TransDecoder, and only those with coding sequences > 90 nucleotides were kept. CD-HIT (Fu et al., 2012) was used to cluster the coding sequences to remove redundancies. Coding sequences of Trinity assembled transcripts were removed if they shared > 90% identity with Iso-Seq isoforms. The remaining coding sequences were mapped to the *P. margaritaceum* genome using GMAP with parameters “-n 0 -z sense force”. According to the alignments, we replaced HC genes predicted from the *P. margaritaceum* genome with corresponding coding transcripts if all the following criteria were met: a) alignment between the transcript and the HC gene covered > 80% of the HC coding region; b) the length of coding sequence of the transcript was at least 1.1 times of that of the HC gene; c) length of the transcript coding sequence should not exceed 1.5 times of the average length of the top 10 protein homologs in the GenBank (www.ncbi.nlm.nih.gov) non-redundant (nr) database (to avoid chimeric transcripts); and d) the transcript was the one that contained longest coding sequence in the locus.

Some transcripts did not align to the genome or the alignments were located in intergenic regions. To identify likely protein-coding genes from among these transcripts, we assessed their coding potential using CPC (Kong et al., 2007) and discarded those annotated as non-coding transcripts. We also excluded the remaining transcripts without any protein homologs in Viridiplantae species, and those whose predicted protein sequences were considered to be too long (≥1.5 times the average length of their homologs) or too short (≤0.5 times the average length of their homologs). In addition, transcripts encoding TE-related Pfam domains were removed. For the remaining transcripts, we only included the longest transcript for each locus into the master gene set.

To annotate genes in the master list, their protein sequences were compared to the GenBank nr database, the *A. thaliana* proteome (www.arabidopsis.org) and the UniProt database (www.uniprot.org) using BLASTp with an E-value cutoff of 1e-5. The protein sequences were also compared to the InterPro database using InterProScan (v5.29-68.0) (Jones et al., 2014). Gene ontology (GO) annotations were obtained using Blast2GO (version 5.2.4) (Gotz et al., 2008) based on the BLASTp results from the GenBank nr database and output from the InterProScan. Functional descriptions were integrated and assigned to the genes using AHRD (v3.3.3; https://github.com/groupschoof/AHRD). Enzyme Commission (EC) information was acquired from the Blast2GO analysis. Transcription factors, transcriptional regulators and protein kinases were identified based on the rules of the iTAK pipeline (v1.7) (Zheng et al., 2016). Pathway analysis was performed using the online annotation server BlastKOALA (Kanehisa et al., 2016).

### Gene family identification

Homologs of each targeted *A. thaliana* gene with known function were identified using a BLASTP E-value ≤ 1e-5 and reciprocal coverage ≥35%. Protein sequences from *P. margaritaceum* and other selected species were used in BLAST searches of the *A. thaliana* protein database (E-value ≤1e-5, coverage of *A. thaliana* genes ≥ 30%), and genes with the best hit to the identified homologs or the original *A. thaliana* genes were identified. Alternatively, some of the gene families were identified based on a search of functional domains with the “--cut-ga” option, or classified based on iTAK (Zheng et al., 2016) results (specified in the Supplementary Table 9).

### Carbohydrate-active enzyme (CAZy) family identification

The CAZY families were identified using dbCAN2 (Zhang et al., 2018), and unless indicated the default thresholds (E-value < 1e-15 and coverage >35%) were used to delineate each gene family.

### Differential gene expression analysis

Cleaned RNA-Seq reads were used to quantify expression of *P. margaritaceum* master genes in each sample using Salmon v0.9.1 (quasi mode; -k 31) (Patro et al., 2017). Raw counts were normalized to FPKM (fragments per kilobase of exon model per million mapped fragments). Differentially expressed genes (DEGs) between treatment and control samples were identified using the DESeq2 package (Love et al., 2014). Genes with false discovery rate (FDR) < 0.01 and fold-change > 2 were considered to be DEGs. GO enrichment analysis of the DEGs was performed using BiNGO (Maere et al., 2005).

### Species phylogeny, molecular dating, and gene family evolution analysis

*P. margaritaceum* and thirteen other species representing major lineages in the taxon Viridiplantae were included for comparative genomics analysis (Supplementary Table 22). The orthogroups among these species were built by OrthoMCL (Li et al., 2003) (v2.0.9) with parameters “E-value < 1e-5; alignment coverage > 40%; inflation value 1.5”. To infer the species phylogeny, we retrieved sequences from low-copy orthogroups, defined as groups in which gene copies for each species was ≤3 and maximum number of species with multi-copy genes, or missing genes, was one. This yielded a total of 738 orthogroups. Each orthogroup was aligned separately with MAFFT and gap regions in the alignment were trimmed with trimAL (Capella-Gutiérrez et al., 2009). A Maximum likelihood phylogeny was inferred by IQ-TREE (Nguyen et al., 2014) (v 1.6.7) with concatenated alignments and best-fitting model (LG+F+I+G4), as well as 1.000 bootstrap replicates. Molecular dating was carried out by MCMCTree in the PAML package (Yang, 2007). The divergence time of Tracheophyta (450.8 - 430.4 Mya) was used as a calibration point according to Morris et al. (2018). For gene family evolution analysis, we used orthogroups with genes present in at least one algal species and one land plant (N=8859). Modeling of gene family size was performed by CAFE (De Bie et al., 2006) (v 4.2) and the gene birth and death rate was estimated with orthogroups that were conserved in all species.

### Whole genome duplication (WGD) analysis

To explore possible WGD in *P. margaritaceum*, we employed CODEML implemented in the PAML package (v4.9h) to obtain *Ks* (synonymous substitution) distribution of paralogous genes. Briefly, *P. margaritaceum* genes in each orthogroup were compared pairwise and gene pairs sharing > 98% identity at both nucleotide and protein levels were eliminated. The *Ks* distribution was fitted with a mixture model of Gaussian distribution by the mclust R package (Scrucca et al., 2016) to identify possible WGD signatures. *Ks* with values >2 were excluded because of Ks saturation. Identification of optimal number of components (corresponding to possible WGDs) in mclust is prone to overfitting, so we also used SiZer and SiCon from the R package (Duong et al., 2008) to distinguish components at a significance level of 0.05.

### Phylogenetic analyses

Protein sequences of the identified genes were aligned using MAFFT (Katoh and Standley, 2013) and the Maximum Likelihood tree for each family was constructed by IQ-TREE (Nguyen et al., 2014) with 1,000 bootstrap replicates. The models used were as specified in the individual tree figures. Abbreviation for selected species (if present) are as follows: *Ostreococcus tauri* (ota, blue), *Chlorella variabilis* (cva, blue), *Chlamydomonas reinhardtii* (cre, blue), *Klebsormidium nitens* (kni, light blue), *Chara braunii* (cbr, light blue), *Penium margaritaceum* (pma, green), *Marchantia polymorpha* (mpo, orange), *Physcomitrella patens* (ppa, orange), *Selaginella moellendorffii* (smo, pink), *Azolla filiculoides* (afi, pink), *Gnetum montanum* (gmo, pink), *Amborella trichopoda* (atr, red), *Oryza sativa* (osa, red) and *Arabidopsis thaliana* (ath, red). Branches with bootstrap value (%) >70 are listed. Orthologs from *P. margaritaceum* and other species were inferred from the tree.

### GH16 sequence characterization and phylogeny construction

BLASTp was used to search through the genomes of *P. margaritaceum, K. nitens* and *C. braunii* with GH16 queries, including laminarinases, agarases, porphyranases, carrageenases, MLGases, chitin transglycosylases, EG16s and XTHs (Viborg et al., 2019). The resulting sequences were analyzed manually and candidates were aligned using SignalP (www.cbs.dtu.dk/services/SignalP), TargetP (www.cbs.dtu.dk/services/TargetP) and a NCBI conserved domain search (www.ncbi.nlm.nih.gov). They were also aligned with other GH16 sequences (using MAFFT, ginsi strategy) and the presence of conserved EG16 and XTH motifs was noted. For the GH16 phylogeny, several representatives of each GH16 group were aligned (MAFFT, ginsi strategy) with all non-fragment *P. margaritaceum* sequences (n=41), *K. nitens* (n=12) sequences, *C. braunii* (n=6) sequences and GH7 cellulases (as outgroup) then trimmed. To calculate the tree, RAxML-HPC2 on XSEDE was used on the CIPRES portal (www.phylo.org), with the JTT amino acid substitution model, which ran for 360 rapid bootstraps.

### Protein structure modeling

The protein structure of *Arabidopsis* ARF1 (PDB accession no. 4LDX) was used as the template for structure modeling of *P. margaritaceum* ARF proteins using SWISS-MODEL (Waterhouse et al., 2018) with default parameters. The target and template sequences were aligned with MAFFT (L-INS-I strategy) and the protein structure was visualized using Chimera (version 1.13.1) (Pettersen et al., 2014).

### Flavonoid extraction and analysis

Flavonoids were extracted from *P. margaritaceum* cell culture pellets based as in Ye et al. (2015). Freeze-dried samples were homogenized with a TissueLyser II (Qiagen), extracted overnight at 4°C with 80% methanol then centrifuged at 13,000 × *g* for 10 min. The supernatant was retained and the pellet re-extracted with 80% methanol for 1 h and centrifuged as before. The supernatants were pooled and dried using a speed-vac then reconstituted in 120 µL of solvent (40 µL MeOH + 80 µL H2O) prior to analysis by liquid chromatography mass spectrometry using a Thermo Scientific Vanquish UHPLC system (mobile phase A, water with 0.1% formic acid; mobile phase B, methanol with 0.1% formic acid), with a Thermo Scientific Hypersil Gold column (2.1 × 150mm, 1.9µm) operating at 45°C with a flow rate of 200 µL min^−1^. Separation of compounds was carried out with gradient elution profile: 0 min, A:B 99.5:0.5; 1 min, A:B 90:10; 10 min, A:B 70:30, 18 min, A:B 50:50, 22 min, A:B 1:99; total 30 min. The injection volume was 2 µL.

MS and MS^n^ data were collected with a Thermo Scientific Orbitrap ID-X Tribrid mass spectrometer. Flavonoids were identified using the automated iterative precursor exclusion method of the Acquire X workflow (four iterative runs of the the pooled sample). The MS^2^ (30 K FWHM at m/z 200) spectra were collected for precursor ions detected in the survey MS scan within a 1.2 second cycle time. Higher order MS^n (3-4)^ (30 K FWHM at m/z 200) spectra were collected only when the instrument detected the sugar neutral loss based on MS^2^ and/or MS^3^ data. For flavonoid quantification, ultra-high resolution MS data (120 K FWHM at m/z 200) was collected. Flavonoid identification and quantification were carried out using Mass Frontier 8.0 and Compound Discoverer 3.1 software (Thermo Scientific). Specifically, flavonoids were identified and annotated searching MS^n^ tree raw data files against mzCloud spectra library using Mass Frontier 8.0. The identified list (full or partial MS^n^ spectral tree data matching MS^n^ spectra of flavonoid references in the mzCloud library) plus an existing database with 6,549 flavonoid structures were used for database search in Compound Discoverer 3.1. The putative flavonoids identified by CD 3.1 were further analyzed for structure annotation by the FISh ranking tool (Mass Frontier 8.0).

### Mucilage compositional analysis

Mucilage was isolated as in Domozych et al. (2005), freeze dried and analyzed by the carbohydrate analytical service of the Complex Carbohydrate Research Center, University of Georgia (www.ccrc.uga.edu/services).

## ACKNOWLEDGMENTS

We thank Sandra Raimundo and Stephen Snyder for technical support and the Imaging, Genomics and facilities of Cornell’s Biotechnology Resource Center, Institute of Biotechnology. J.K.C.R. is supported by the Cornell Atkinson Center for Sustainability. The mucilage analysis work was supported by the Chemical Sciences, Geosciences and Biosciences Division, Office of Basic Energy Sciences, U.S. Department of Energy grant (DE-SC0015662) to Dr. Parastoo Azadi at the Complex Carbohydrate Research Center, University of Georgia.

## AUTHOR INFORMATION

The authors declare no competing interests

## REFERENCES

Agarwal, P.K., Gupta, K., Lopato, S. & Agarwal, P. Dehydration responsive element binding transcription factors and their applications for the engineering of stress tolerance. J. Exp. Bot. 68, 2135–2148 (2017).

Bak, S. et al. Cytochromes p450. The arabidopsis book 9, e0144–e0144 (2011).

Bankevich, A. et al. SPAdes: a new genome assembly algorithm and its applications to single-cell sequencing. J. Comput. Biol. 19, 455–477 (2012).

Bateman, A. et al. The Pfam protein families database. Nucleic Acids Res. 32, D138–D141 (2004).

Behar, H., Graham, S.W. & Brumer, H. Comprehensive cross-genome survey and phylogeny of glycoside hydrolase family 16 members reveals the evolutionary origin of EGE16 and XTH proteins in plant lineages. Plant J. 95, 1114–1128 (2018).

Benson, G. Tandem repeats finder: a program to analyze DNA sequences. Nucleic Acids Res. 27, 573–580 (1999).

Bolger, A.M., Lohse, M. & Usadel, B. Trimmomatic: a flexible trimmer for Illumina sequence data. Bioinformatics 30, 2114–2120 (2014).

Bolle, C. The role of GRAS proteins in plant signal transduction and development. Planta 218, 683–692 (2004).

Boney, A.D. Mucilage: a ubiquitous algal attributer. Brit. Phyco. J. 16, 115–132 (1981).

Boot, K.J.M., Libbenga, K.R., Hille, S.C., Offringa, R. & van Duijn, B. Polar auxin transport: an early invention. J. Exp. Bot. 63, 4213–4218 (2012).

Bowman, J.L., Briginshaw, L.N., Fisher, T.J. & Flores-Sandoval, E. Something ancient and something neofunctionalized—evolution of land plant hormone signaling pathways. Curr. Opin. Plant Biol. 47, 64–72 (2019).

Brook, A. The biology of desmids. Berkeley: University of California Press (1981).

Burton, R. A., Gidley, M. J. & Fincher, G. B. Heterogeneity in the chemistry, structure and function of plant cell walls. *Nature Chem*. Biol. 6, 724–732 (2010).

Campbell, M.S. et al. MAKER-P: a tool kit for the rapid creation, management, and quality control of plant genome annotations. Plant Physiol. 164, 513–524 (2014).

Cantarel, B.L. et al. The Carbohydrate-Active EnZymes database (CAZy): an expert resource for glycogenomics. Nucl. Acids Res. 37, D233–D238 (2009).

Cantarel, B.L. et al. MAKER: an easy-to-use annotation pipeline designed for emerging model organism genomes. Genome research 18, 188–196 (2008).

Capella-Gutiérrez, S., Silla-Martínez, J.M. & Gabaldón, T. trimAl: a tool for automated alignment trimming in large-scale phylogenetic analyses. Bioinformatics 25, 1972–1973 (2009).

Catarino, B., Hetherington, A.J., Emms, D.M., Kelly, S. & Dolan, L. The stepwise increase in the number of transcription factor families in the Precambrian predated the diversification of plants on land. Mol. Biol. Evol. 33, 2815–2819 (2016).

Chin, C.-S. et al. Nonhybrid, finished microbial genome assemblies from long-read SMRT sequencing data. Nature Methods 10, 563 (2013).

Clavijo, B.J. et al. An improved assembly and annotation of the allohexaploid wheat genome identifies complete families of agronomic genes and provides genomic evidence for chromosomal translocations. Genome Research 27, 885–896 (2017).

Cosgrove, D.J. Re-constructing out models of cellulose and primary cell wall assembly. Curr. Opin. Plant Biol. 22, 122–131 (2014).

Cosgrove, D.J. Plant expansins: diversity and interactions with plant cell walls. Curr. Opin. Plant Biol. 25, 162–172 (2015).

Cutler, S.R., Rodriguez, P.L., Finkelstein, R.R. & Abrams, S.R. Abscisic acid: emergence of a core signaling network. Annu. Rev. Plant Biol. 61, 651–679 (2010).

De Bie, T., Cristianini, N., Demuth, J.P. & Hahn, M.W. CAFE: a computational tool for the study of gene family evolution. Bioinformatics 22, 1269–1271 (2006).

deVries, J., Curtis, B.A., Gould, S.B. & Archibald, J.M. Embryophyte stress signaling evolved in algal progenitors of land plants. Proc. Natl. Acad. Sci. USA. 115, E3471–3480 (2018).

de Vries, J., de Vries, S., Slamovits, C.H., Rose, L.E. & Archibald, J.M. How embryophytic is the biosynthsis of phenylpropanoids and their derivatives in streptophyte algae? Plant Cell Physiol. 58, 934–945 (2017).

Delaux, P.-M. et al. Algal ancestor of land plants was preadapted for symbiosis. Proc. Natl. Acad. Sci. USA 112, 13390–13395 (2015).

Delwiche, C.F. & Cooper, E.D. The evolutionary origin of a terrestrial flora. Curr. Biol. 25, R889–R910 (2015).

Delwiche, C.F. & Timme, R.E. Plants. Curr. Biol. 21, R417–422 (2011).

Dehors, J. et al. Evolution of cell wall polymers in tip-growing land plant gametophytes: composition, distribution, functional aspects and their remodeling. Front. Plant Sci. 10, 441 (2019).

Domozych, D.S. & Domozych, C.R. Desmids and biofilms of freshwater wetlands: development and microarchitecture. Microbial Ecol. 55, 81–93 (2008).

Domozych, D.S., Kort, S., Benton, S. & Yu, T. The extracellular polymeric substance of the green alga *Penium margaritaceum* and its role in biofilm formation. Biofilms 2, 129–144 (2005).

Domozych, D.S., Serfis, A., Kiemle, S.N. & Gretz, M.R. The structure and function of charophycean cell walls. I Pectins of *Penium margaritaceum*. Protoplasma 239, 99–115 (2007).

Domozych, D.S. et al. Pectin metabolism and assembly in the cell wall of the charophyte green alga *Penium margaritaceum*. Plant Physiol. 165, 105–118 (2014).

Duong, T., Cowling, A., Koch, I. & Wand, M.P. Feature significance for multivariate kernel density estimation. Comput. Stat. Data Anal. 52, 4225–4242 (2008).

Eklund, D.M. et al. An evolutionarily conserved abscisic acid signaling pathway regulates dormancy in the liverwort *Marchantia polymorpha*. Curr. Biol. 28, 36912–3699 (2019).

Ellinghaus, D., Kurtz, S. & Willhoeft, U. LTRharvest, an efficient and flexible software for de novo detection of LTR retrotransposons. BMC Bioinformatics 9, 18 (2008).

Elköf, J.M., Shojania, S., Okon, M., McIntosh, L.P. & Brumer, H. Structure-function analysis of a broad specificity *Populus trichocarpa* endo-β-glucanase reveals an evolutionary link between bacterial licheninases and plant XTH gene products. J. Biol. Chem. 288, 15786–15799 (2013).

Emiliani, G., Fondi, M., Fani, R., & Gribaldo, S. A horizontal gene transfer at the origin of phenylpropanoid metabolism: a key adaptation of plants to land. Biol. Direct. 4, 7 (2009).

Engstrom, E.M. Phylogenetic analysis of GRAS proteins from moss, lycophyte and vascular plant lineages reveals that GRAS genes arose and underwent substantial diversification in the ancestral lineage common to bryophytes and vascular plants. Plant signaling & behavior 6, 850–854 (2011).

Feist, M., Liu, J. & Tafforeau, P. New insights into Paleozoic charophyte morphology and phylogeny. Am. J. Bot. 92, 1152–1160 (2005).

Fich, E.A., Segerson, N.A. & Rose, J.K.C. The plant polyester cutin: biosynthesis, structure and biological roles. Ann. Rev. Plant Biol. 67, 207–233 (2016).

Fisher, M.M., Wilcox, L.W. & Graham, L.E. Molecular characterization of epiphytic bacterial communities on charophycean green algae. Appl. Environ. Microbiol 64, 4384–4389 (1998).

Fu, L., Niu, B., Zhu, Z., Wu, S. & Li, W. CD-HIT: accelerated for clustering the next-generation sequencing data. Bioinformatics 28, 3150–3152 (2012).

Garrison, E. & Marth, G. Haplotype-based variant detection from short-read sequencing. arXiv preprint arXiv, 12073907 (2012).

Gnerre, S. et al. High-quality draft assemblies of mammalian genomes from massively parallel sequence data. Proc. Natl. Acad. Sci. USA 108, 1513–1518 (2011).

Gotz, S. et al. High-throughput functional annotation and data mining with the Blast2GO suite. Nucleic Acids Res. 36, 3420–3435 (2008).

Grabherr, M.G. et al. Full-length transcriptome assembly from RNA-Seq data without a reference genome. Nat. Biotechnol. 29, 644–652 (2011).

Haas, B.J. et al. Improving the Arabidopsis genome annotation using maximal transcript alignment assemblies. Nucleic Acids Res. 31, 5654–5666 (2003).

Han, Y. & Wessler, S.R. MITE-Hunter: a program for discovering miniature inverted-repeat transposable elements from genomic sequences. Nucleic Acids Res. 38, e199 (2010).

Hauser, F., Waadt, R. & Schroeder, J. I. Evolution of abscisic acid synthesis and signaling mechanisms. Curr. Biol. 21, R346–R355 (2011).

Herburger, K. & Holzinger, A. Localization and quantification of callose in the streptophyte green algae *Zygnema* and *Klebsormidium*: correlation with desiccation tolerance, Plant Cell Phys. 56, 2259–2270 (2015).

Hernandez-Garcia, J., Briones-Moreno, A., Dumas, R., & Blazquez, M.A. Origin of gibberellin-dependent transcriptional regulation by molecular exploitation of a transactivation domain in DELLA proteins. Mol. Biol. Evol. 36,: 908–918 (2019).

Holzinger, A. & Pichrtova, M. Abiotic stress tolerance of charophyte green algae: new challenges for omics techniques. Front. Plant Sci. 7, 678 (2016).

Hong, L., Brown, J., Segerson, N., Rose, J.K.C. & Roeder, A.H.K. CUTIN SYNTHASE2 maintains progressively developing cuticular ridges in *Arabidopsis* sepals. Mol. Plant 10, 560–574 (2017).

Hori, K. et al. *Klebsormidium flaccidum* genome reveals primary factors for plant terrestrial adaptation. Nat Commun 5, 3978 (2014).

Jackman, S.D. et al. ABySS 2.0: resource-efficient assembly of large genomes using a Bloom filter. Genome research 27, 768–777 (2017).

Jones, P. et al. InterProScan 5: genome-scale protein function classification. Bioinformatics 30, 1236–1240 (2014).

Ju, C. et al. Conservation of ethylene as a plant hormone over 450 million years of evolution. Nat Plants 1, 14004 (2015).

Kanehisa, M., Sato, Y. & Morishima, K. BlastKOALA and GhostKOALA: KEGG Tools for Functional Characterization of Genome and Metagenome Sequences. J. Mol. Biol. 428, 726–731 (2016).

Katoh, K. & Standley, D.M. MAFFT multiple sequence alignment software version 7: improvements in performance and usability. Mol. Biol. Evol. 30, 772–780 (2013).

Kazmierczak, A. & Rosiak, M. Content of gibberellic acid in apical parts of male and female thalli of *Chara tomentosa* in relation to the content of sugars and dry mass. Biologia Plantarum 43, 369–372 (2000).

Kiemle, S.N., Domozych, D.S. & Gretz, M.R. The extracellular polymeric substances of desmids (Conjugatophyceae, Streptophyta): chemistry, structural analyses and implications in wetland biofilms. Phycologia 46, 617–627 (2007).

Kim, D., Langmead, B. & Salzberg, S.L. HISAT: a fast spliced aligner with low memory requirements. Nature Methods 12, 357 (2015).

Kim, T.W. et al. Brassinosteroid signal transduction from cell-surface receptor kinases to nuclear transcription factors. Nat Cell Biol 11, 1254–1260 (2009).

Kiseleva, A.A., Tarachovskaya, E.R., & Shisova, M.F. Biosynthesis of phytohormones in algae. Russ. J. Plant Physiol. 59, 595–610 (2012).

Knox, J.P., Linstead, P.J., King, J., Cooper, C. & Roberts, K. Pectin esterification is spatially regulated both within cell walls. Planta, 181, 512–521 (1990).

Kollist, H. et al. Rapid responses to abiotic stress: priming the landscape for the signal transduction network. Trends Plant Sci. 24, 25–37 (2019).

Kondo, S. et al. Primitive extreacellular lipid components on the surface of the charophytic alga *Klebsormidium flaccidum* and their possible biosynthetic pathways as deduced from the genome sequence. Frontiers Plant Sci. 7, 952 (2016).

Kong, L. et al. CPC: assess the protein-coding potential of transcripts using sequence features and support vector machine. Nucleic Acids Res. 35, W345–349 (2007).

Korf, I. Gene finding in novel genomes. BMC Bioinformatics 5, 59 (2004).

Kurakawa, T. et al. Direct control of shoot meristem activity by a cytokinin-activating enzyme. Nature 445, 652 (2007).

Kuromori, T., Seo, M. & Shinozaki, K. ABA transport and plant water stress reponses. Trends Plant Sci. 23, 513–522 (2018).

Lang, D. et al. Genome-wide phylogenetic comparative analysis of plant transcriptional regulation: a timeline of loss, gain, expansion, and correlation with complexity. Genome Biol Evol 2, 488–503 (2010).

Lashbrooke, J. et al. MYB17 and MYB9 homologs regulate suberin deposition in angiosperms. Plant Cell 28, 2097–2116 (2016).

Lee, S.B. & Suh, M.C. Advances in the undertanding of cuticular waxes in *Arabidopsis thaliana* and crop species. Plant Cell Reports 34, 557–572 (2015).

Leebens-Mack, J.H. et al. One thousand plant transcriptomes and the phylogenomics of green plants. Nature, doi:10.1038/s41586-019-1693-2 (2019).

Leyser, O. Auxin signaling. Plant Physiol. 176, 465–479 (2018).

Li, D., Liu, C.M., Luo, R., Sadakane, K. & Lam, T.W. MEGAHIT: an ultra-fast single-node solution for large and complex metagenomics assembly via succinct de Bruijn graph. Bioinformatics 31, 1674–1676 (2015).

Li, H. Minimap2: pairwise alignment for nucleotide sequences. Bioinformatics 34, 3094–3100 (2018).

Li, H. & Durbin, R. Fast and accurate short read alignment with Burrows-Wheeler transform. Bioinformatics 25, 1754–1760 (2009).

Li, L., Stoeckert, C.J. & Roos, D.S. OrthoMCL: identification of ortholog groups for eukaryotic genomes. Genome Res 13, 2178–2189 (2003).

Li, Z., Shen, J., & Liang, J. Genome-wide identification, expression profile, and alternative splicing anlaysis of the brassinosteroid-signaling kinase (BSK) family genes in *Arabidopsis*. Int. J. Mol. Sci. 20, 1138 (2019).

Lievens, L., Pollier, J., Gooseens, A., Beyaert, R., & Staal, J. Abscisic acid as pathogen effector and immune regulator. Frontiers Plant Sci. 8, 587 (2017).

Lind, C. et al. Stomatal guard cells co-opted an ancient ABA-dependent desiccation survival system to regulate stomatal closure. Curr. Biol. 25, 928–935 (2015).

Lombard, V. et al. A heirarchical classification of polysacchaide lyases for glycogenomics. Biochem. J. 432, 437–444 (2010).

Lomsadze, A., Ter-Hovhannisyan, V., Chernoff, Y.O. & Borodovsky, M. Gene identification in novel eukaryotic genomes by self-training algorithm. Nucleic acids res. 33, 6494–6506 (2005).

Love, M.I., Huber, W. & Anders, S. Moderated estimation of fold change and dispersion for RNA-seq data with DESeq2. Genome Biol 15, 550 (2014).

Lu, Y. & Xu, J. Phytohormones in microalgae: a new opportunity for microalgal biotechnology? Trends Plant Sci. 20, 273–282 (2015).

Luo, R. et al. SOAPdenovo2: an empirically improved memory-efficient short-read de novo assembler. Gigascience 1, 18 (2012).

Ma, H.S., Liang, D., Shuai, P., Xia, X.L. & Yin, W.L. The salt- and drought-inducible poplar GRAS protein SCL7 confers salt and drought tolerance in *Arabidopsis thaliana*. J. Exp. Bot. 61, 4011–4019 (2010).

Maere, S., Heymans, K. & Kuiper, M. BiNGO: a Cytoscape plugin to assess overrepresentation of gene ontology categories in biological networks. Bioinformatics 21, 3448–3449 (2005).

Mapleson, D., Garcia Accinelli, G., Kettleborough, G., Wright, J. & Clavijo, B.J. KAT: a K-mer analysis toolkit to quality control NGS datasets and genome assemblies. Bioinformatics 33, 574–576 (2016).

Martin-Arevalillo, R. et al. Evolution of the auxin response factors from charophyte ancestors. PLOS Genet 15, e1008400 (2019).

McGregor, N., Yin, V., Tung, C.C., Van Petergem, F., & Brumer, H. Crystallographic insight into the evolutionary origins of xyloglucan endotransglycosylases and endohydrolases. Plant J. 89, 651–670 (2017).

Morgan, M. et al. ShortRead: a bioconductor package for input, quality assessment and exploration of high-throughput sequence data. Bioinformatics 25, 2607 (2009).

Morris, J.L. et al. The timescale of early land plant evolution. Proc. Natl. Acad. Sci. USA 115, E2274–e2283 (2018).

Nadalin, F., Vezzi, F. & Policriti, A. GapFiller: a de novo assembly approach to fill the gap within paired reads. BMC Bioinformatics 13 Suppl 14, S8 (2012).

Narasimhan, V. et al. BCFtools/RoH: a hidden Markov model approach for detecting autozygosity from next-generation sequencing data. Bioinformatics 32, 1749–1751 (2016).

Nguyen, L.-T., Schmidt, H.A., von Haeseler, A. & Minh, B.Q. IQ-TREE: a fast and effective stochastic algorithm for estimating maximum-likelihood phylogenies. Mol. Biol. Evol. 32, 268–274 (2014).

Nichols, H.W. Growth media-freshwater. In: Stein JR, editor. Handbook of phycological methods: culture methods and growth measurements. New York: Cambridge University Press; 1973. p. 39–78.

Niklas, K.J., Cobb, E.D. & Matas, A.J. The evolution of hydrophobic cell wall biopolymers: from algae to angiosperms. J. Exp. Bot. 68, 5261–5269 (2017).

Nishiyama, T. et al. The *Chara* genome: secondary complexity and implications for plant terrestrialization. Cell 174, 448–464. e424 (2018).

Oertel, A., Aichinger, N., Hochreiter, R., Thalhammer, J. & Lütz-Meindl, U. Analysis of mucilage secretion and excretion in *Micrasterias* (Chlorophyta) by means of immunoelectron microscopy and digital time lapse video microscopy. J. Phycol. 40, 711–720 (2004).

Ohtaka, K., Hori, K., Kanno, Y., Seo, M. & Ohta, H. Primitive auxin response without TIR1 and Aux/IAA in the charophyte alga *Klebsormidium nitens*. Plant Physiol. 174, 1621–1632 (2017).

Ossowski, S. et al. The rate and molecular spectrum of spontaneous mutations in *Arabidopsis thaliana*. Science 327, 92–99 (2010).

Panchy, N., Lehti-Shiu, M. & Shiu, S.-H. Evolution of gene duplication in plants. Plant Physiol. 171, 2294–2316 (2016).

Pascal, S. et al. Arabidopsis CER1-LIKE1 functions in a cuticular very-long-chain alkane-forming complex. Plant Physiol. 179, 415–432 (2019).

Patro, R., Duggal, G., Love, M.I., Irizarry, R.A. & Kingsford, C. Salmon provides fast and bias-aware quantification of transcript expression. Nature Methods 14, 417 (2017).

Pertea, M. et al. StringTie enables improved reconstruction of a transcriptome from RNA-seq reads. Nature Biotechnology 33, 290 (2015).

Pettersen, E.F. et al. UCSF Chimera—a visualization system for exploratory research and analysis. J. Comp. Chem. 25, 1605–1612 (2004).

Pineau, E. et al. *Arabidopsis thaliana* EPOXIDE HYDROLASE1 (AtEH1) is a cytosolic epoxide hydrolase involved in the synthesis of poly-hydroxylated cutin monomers. New Phytol. 215, 173–186 (2017).

Popper, Z.A. et al. Evolution of diversity of plant cell walls: from algae to flowering plants. Ann. Rev. Plant Biol. 62, 567–590 (2011).

Quast, C. et al. The SILVA ribosomal RNA gene database project: improved data processing and web-based tools. Nucleic Acids Res. 41, D590–596 (2013).

Raimundo, S.C. et al. Protoplast isolation and manipulation of protoplasts from the unicellular green alga *Penium margaritaceum*. Plant Methods 14, 18 (2018).

Ren, H. et al. BRASSINOSTEROID-SIGNALING KINASE 3, a plasma membrane-associated scaffold protein involved in early brassinosteroid signling. PLoS Genet. 15, e1007904 (2019).

Renault, H., Werck-Reichhart, D. & Weng, J.-K. Harneessing lignin evolution for biotechnological applications. Curr. Opin. Plant Biotechnol. 56, 105–111 (2019).

Rensing, S.A. Great moments in evoluiton: the conquest of land by plants. Curr. Opin. Plant Biol. 42, 49–54 (2018).

Rice, P., Longden, I. & Bleasby, A. EMBOSS: the European molecular biology open software suite. Trends in genetics 16, 276–277 (2000).

Romani, F. Origin of TAA genes in Charophytes: New insights into the controversy over the origin of auxin biosynthesis. Frontiers in Plant Science 8, 1616 (2017).

Rossel, J.B. et al. Systemic and intracellular responses to photooxidative stress in Arabidopsis. Plant Cell 19, 4091–4110 (2007).

Rydahl, M.G. et al. *Penium margaritaceum* as a model organism for cell wall analysis of expanding plant cells. In: Estevez, J.M., ed., Plant Cell Expansion. Methods in Molecular Biology. New York, Springer 1242, 1–21 (2015).

Salmela, L. & Rivals, E. LoRDEC: accurate and efficient long read error correction. Bioinformatics 30, 3506–3514 (2014).

Scrucca, L., Fop, M., Murphy, T.B. & Raftery, A.E. mclust 5: Clustering, Classification and Density Estimation Using Gaussian Finite Mixture Models. The R journal 8, 289–317 (2016).

Shanmugarajah, K. et al. ABCG1 contributes to suberin formation in *Arabidopsis thaliana* roots. Scientific Reports 9, 11381 (2019).

Siewers, V., Kokkelink, L., Smedsgaard, J. & Tudzynski, P. Identification of an abscisic acid gene cluster in the grey mold Botrytis cinerea. Appl. environ. microb. 72, 4619–4626 (2006).

Signorelli, S., Tarkowski, L.P., Van den Ende, W. & Bassham, D.C. Linking autophagy to abiotic and biotic stress repsonses. Trends Plant Sci. 24, 413–430 (2019).

Simão, F.A., Waterhouse, R.M., Ioannidis, P., Kriventseva, E.V., Zdobnov, E.M. BUSCO: assessing genome assembly and annotation completeness with single-copy orthologs. Bioinformatics 31, 3210–3212 (2015).

Sørensen, I. et al. Stable transformation and reverse genetic analysis of *Penium margaritaceum*: a platform for studies of charophyte green algae, the immediate ancestors of land plants. Plant J. 77, 339–351 (2014).

Sørensen, I. et al. The charophycean green algae provide insights into the early origins of plant cell walls. Plant J. 68, 201–211 (2011).

Stanke, M., Tzvetkova, A. & Morgenstern, B. AUGUSTUS at EGASP: using EST, protein and genomic alignments for improved gene prediction in the human genome. Genome Biol 7 Suppl 1, S11.1–8 (2006).

Swarup, R. & Bennett, M.J. Auxin transport:providing plants with a new scale new sense of direction. Biochem. Soc. Trans. 36, 12–16 (2014).

Swarup, R. & Bhosale, R. Developmental roles of AUX1/LAX auxin influx carrieres in plants. Frontiers Plant Sci., https://doi.org/10.3389/fpls.2019.01306 (2019).

Sun, Y. et al. Integration of brassinostroid signal transduction with the transcription network for plant growth regulation in *Arabidopsis*. Dev. Cell 19, 765–777 (2010).

Suzek, B.E. et al. UniRef clusters: a comprehensive and scalable alternative for improving sequence similarity searches. Bioinformatics 31, 926–932 (2014).

Tenhaken, R. Cell wall remodeling under abiotic stress. Frontiers Plant Sci. 5, 771 (2015).

Terrett, O. & Dupree, P. Covalent interactions between lignin and hemicelluloses in plant secondary cell walls. Curr. Opin. Plant Biotechnol. 56, 97–104 (2019).

Thalmann, M. & Santelia, D. Starch as a deterinant of plant fitness under abiotic stress. New Phytol. 214, 943–951 (2017).

Tivendale, N.D., Ross, J.J. & Cohen, J.D. The shifting paradigms of auxin biosynthesis. Trends Plant Sci. 19, 44–51 (2014).

Van de Peer, Y., Mizrachi, E. & Marchal, K. The evolutionary significance of polyploidy. Nat. Rev. Genet. 18, 411–424 (2017).

Van de Poel, B., Cooper, E.M., Van der Straeten, D., Chang C. and Delwiche, C.F. Transcriptome profiling of the green alga *Spirogyra pratense* (Charophyta) suggests an ancestral role for ethylene in cell wall meabolism, photosynthesis and abiotic stress responses. Plant Physiol. 172, 533–545 (2016).

Viaene, T., Delwiche, C.F., Rensing, S.A. & Friml, J. Origin and evolution of PIN auxin transporters in the green lineage. Trends Plant Sci. 18, 5–10 (2012).

Viborg, A.H. et al. A subfamily roadmap of the evolutionarily diverse glycoside hydrolase family 16 (GH16). J. Biol. Chem. 294, 15973–15986 (2019).

Vishwanath, S.J., Delude, C., Domergue, F. & Rowland, O. Suberin: biosyntehsis, regulation, and polymer assembly of a protective exatrcellular barrier. Plant Cell Reports 34, 573–586 (2015).

Walker, C.H., Siu-Ting, K., Taylore, A., O’Connell, M.J. & Bennett, T. Strigolactone synthesis is ancestral in land plants, but canonical strigolactone signalling is a flowering plant innovation. BMC Biol. 17, 70 (2019).

Wan, C.Y. & Wilkins, T.A. A modified hot borate method significantly enhances the yield of high-quality RNA from cotton. Anal. Biochem. 223, 7–12 (1994).

Waterhouse, A. et al. SWISS-MODEL: homology modelling of protein structures and complexes. Nucleic Acids Res. 46, W296–W303 (2018).

Weng, J.-K. and Chapple, C. The origin and evolution of lignin biosynthesis. New Phytol. 187, 273–285 (2010).

Wodniok, S. et al. Origin of land plants: do conjugating green algae hold the key? BMC Evol. Biol. 11, 104 (2011).

Wu, T.D. & Watanabe, C.K. GMAP: a genomic mapping and alignment program for mRNA and EST sequences. Bioinformatics 21, 1859–1875 (2005).

Yang, Z. PAML 4: phylogenetic analysis by maximum likelihood. Mol. Biol. Evol. 24, 1586–1591 (2007).

Ye, J. et al. Transcriptome profiling of tomato fruit development reveals transcription factors associated with ascorbic acid, carotenoid and flavonoid biosynthesis. PLOS ONE 10, e0130885 (2015).

Yeats, T.H. & Rose, J.K.C. The formation and function of plant cuticles. Plant Physiol. 163, 5–20 (2013).

Yonekura-Sakakibara, K., Higashi, Y. & Nakabayashi, R. The origin and evolution of plant flavonoid metabolism. Frontiers Plant Sci. 10, 943 (2019).

Yue, J., Hu, X., Sun, H., Yang, Y. & Huang, J. Widespread impact of horizontal gene transfer on plant colonization of land. Nat Commun 3, 1152 (2012).

Żabka, A. et al. PIN2-like proteins may contribute to the regulation of morphogenetic processes during spermatogenesis in *Chara vulgaris*. Plant Cell Reports 35, 1655–1669 (2016).

Zamil, M.S. & Geitmann, A. The niddle lamella-more than a glue. Phys. Biol. 14, 015004 (2017).

Zhang, D., Iyer, L.M. & Aravind, L. Bacterial GRAS domain proteins throw new light on gibberellic acid response mechanisms. Bioinformatics 28, 2407–2411 (2012).

Zhang, H. et al. dbCAN2: a meta server for automated carbohydrate-active enzyme annotation. Nucleic Acids Res. 46, W95–W101 (2018).

Zheng, Y. et al. iTAK: A program for genome-wide prediction and classification of plant transcription factors, transcriptional regulators, and protein kinases. Mol. Plant 9, 1667–1670 (2016).

Zheng, Y., Zhao, L., Gao, J. & Fei, Z. iAssembler: a package for de novo assembly of Roche-454/Sanger transcriptome sequences. BMC Bioinformatics 12, 453 (2011).

Zhong, R., Cui, D. & Ye, Z.-H. Secondary cell wall biosynthesis. New Phytol. 221, 1703–1723 (2019).

Zhong, S. et al. High-throughput illumina strand-specific RNA sequencing library preparation. Cold Spring Harb Protoc 2011, 940–949 (2011).

Zimin, A.V. et al. The MaSuRCA genome assembler. Bioinformatics 29, 2669–2677 (2013).

